# Repetitive anatomical patterns for thalamocortical projections of higher-order thalamic nuclei

**DOI:** 10.64898/2026.06.25.734453

**Authors:** Andreas Huth, Thomas Kuner

## Abstract

Cortico-thalamo-cortical circuits entail extensive trans-thalamic connectivity between cortical areas, yet their structural organization and function remain poorly understood. Here, the thalamocortical projections of several higher-order thalamic nuclei were characterized by retrograde tracing from two cortical areas, the primary somatosensory (S1) and motor (M1) cortices. Cholera toxin B conjugated with different fluorophores allowed for simultaneous detection of projection neurons targeting S1 and M1. A cell detection pipeline based on neural networks was developed to allow semi-automated analysis of large thalamic imaging volumes to quantitatively infer the spatial distribution of projection neurons in the posterior complex (PO) and the adjacent ethmoid nucleus (Eth), nucleus centrolateralis (CL), nucleus paracentralis (PCN), and the nucleus parafascicularis (PF). The arrangement of neurons projecting to both, primary somatosensory and motor cortices, occurs at different connection strengths and was topographically organized in all nuclei studied. Co-injections into both cortical areas revealed projection neurons with axons branching into both S1 and M1 cortices. Our work introduces a pipeline for semi-automated quantitative analysis of thalamic projection patterns that could be useful for connectivity analyses in general. This approach revealed repetitive anatomical patterns in different thalamic nuclei with regard to projection strength, spatial organization and fraction of projection neurons targeting two cortical areas simultaneously.

## Introduction

Thalamic nuclei can be classified as first-order and higher-order thalamic nuclei (Guillery, 1995; Sherman & Guillery, 2002). First-order thalamic nuclei are a relay station for peripheral information on the way to the cortex (Sherman & Guillery, 2002). In contrast, higher-order thalamic nuclei appear to have an important role in cortico-thalamocortical (CTC) information processing (Blot et al., 2021; Sherman & Guillery, 2002). Thus, CTC connections represent a second, parallel route of cortico-cortical information transfer compared to the direct transcortical pathway (Sherman, 2016; Sherman & Guillery, 2011). The exact function of this trans-thalamic route is largely unknown (Theyel et al., 2010). However, different information is transmitted to cortical areas via the trans-thalamic route compared to the more direct cortico-cortical route (Blot et al., 2021; Roth et al., 2016). For example, CTC connections may accentuate information in cortical areas (Mease et al., 2016) and synchronize cortical areas (Saalmann, 2014; Saalmann et al., 2012).

The projections of the posterior complex of the thalamus (PO) have been shown to target the primary (S1) and secondary (S2) somatosensory cortex, primary (M1) and secondary (M2) motor cortex, and other cortical areas such as the insular cortex (Casas-Torremocha et al., 2022; Miller-Hansen & Sherman, 2022; Oh et al., 2014; Sampathkumar et al., 2021). However, the precise topographical organization and existence of PO axons targeting several target regions at once remained poorly characterized. The same holds for the adjacent ethmoid nucleus (Eth), the nucleus centrolateralis (CL), the nucleus paracentralis (PCN), and the nucleus parafascicularis (PF). The latter three belong to the intralaminar nuclei, which may contribute to consciousness and arousal (Vertes et al., 2022). The PO, as it is defined in the Allen Mouse Brain Common Coordinate Framework in the Version 3 (CCFv3) (Wang et al., 2020), is in parts identical to the posteromedial thalamic nucleus (POm) (see discussion for more details). POm, Eth and the intralaminar nuclei can be classified as higher-order thalamic nuclei (Bokor et al., 2005; Mease & Gonzalez, 2021; Saalmann, 2014; Van Horn & Sherman, 2007).

In this study, retrograde tracing was performed in mice to further investigate thalamocortical projections. A cell detection pipeline has been developed using cell positions in approximately 17 consecutive brain slices per animal. Thus, cell positions could be reconstructed for a volume of 2–3 mm^3^ of the right thalamus and individual cells were assigned to the corresponding thalamic nuclei. Subsequent analysis revealed general patterns in the thalamic nuclei with respect to projection strength, topographic arrangement of cells, and potential substructures in the thalamic nuclei. Co-injections into both cortical areas showed that some projection neurons target S1 and M1 simultaneously.

## Results

Neurons within each thalamic nucleus were labeled by retrograde tracing using Cholera toxin B conjugated with the fluorescent dyes Alexa Fluor 488, Alexa Fluor 555, or Alexa Fluor 647 (CTB-Af488, CTB-Af555 or CTB-Af647) injected into S1 or M1 (Fig. 1a, Supp. Tab. 1–3). The three experimental groups consisted of injections only in S1 (13 injections in five mice), injections only in M1 (9 injections in five mice), and co-injections in S1 and M1 (12 injections in six mice). After perfusion of the animals and removal of the brains, overview images of the brain slices were taken using a wide-field fluorescence microscope and image stacks with a higher resolution were acquired using laser scanning confocal microscopy (LSCM). No crosstalk between the channels was detected for LSCM (Supp. Fig. 1). Positions of the injection sites were obtained by adapting brain slices containing the signal of the injection site to corresponding reference atlas slices (Fig. 1b, Supp. Fig. 2).

**Figure 1.**
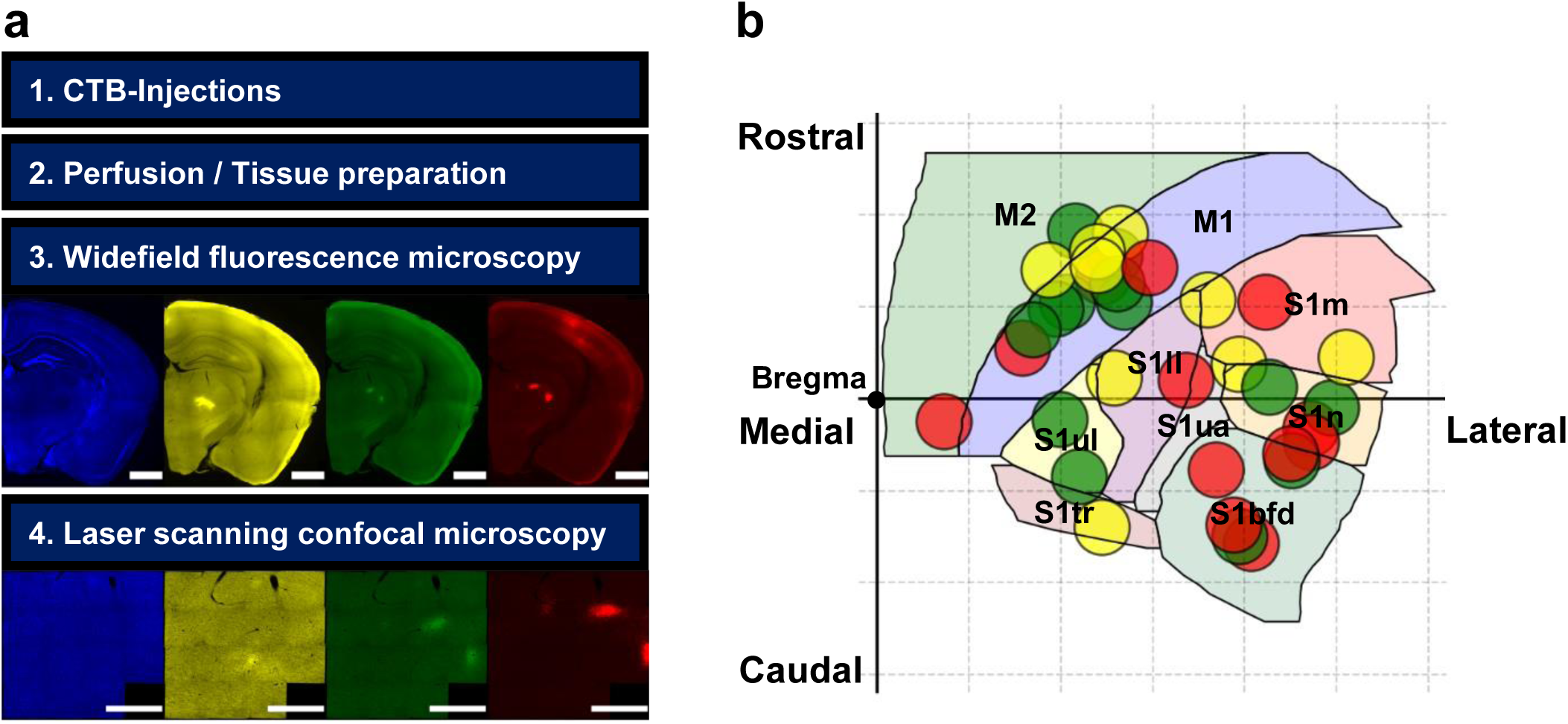
Experimental procedure. (a) General experimental procedure. Blue: DAPI-channel, yellow: CTB-Af488-channel, green: CTB-Af555-channel, red: CTB-Af647-channel. Scale bar: 1 mm (upper four images) and 0.5 mm (lower four images). (b) Position of the injection sites in the primary somatosensory cortex (*n* = 19 injections in 11 mice) and the primary motor cortex (*n* = 15 injections in 11 mice). Yellow: CTB-Af488 injections, green: CTB-Af555 injections, red: CTB-Af647 injections. The grid spacing is 1 mm. M1: primary motor cortex, M2: secondary motor cortex, S1: primary somatosensory cortex with several subareas (mouth area (S1m), nose area (S1n), barrel field (S1bfd), lower limb area (S1ll), upper limb area (S1ul), trunk area (S1tr) and unassigned area (S1ua)). Alt text: Graphical representations of the experimental procedure.

**Figure 2.**
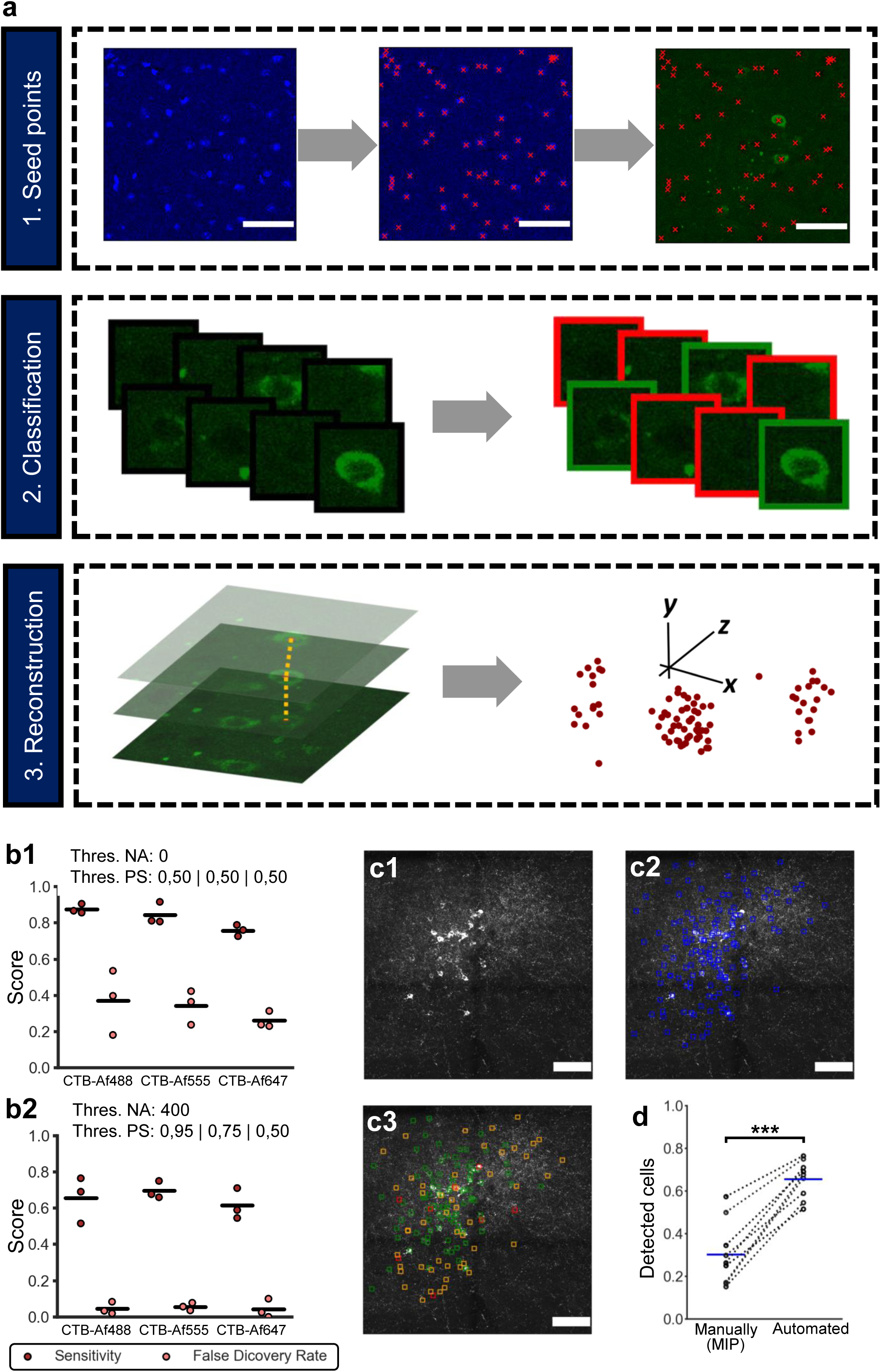
Cell detection pipeline. (a) Steps of the cell detection. In a first step the positions of the cell nuclei (red crosses) are detected (top). This information is used to obtain images which can then be classified whether they represent a labeled cell (green frame) or not (red frame) (middle). Detections of the same cell in multiple layers are identified (indicated by dashed orange lines) to avoid multiple registrations. Positions of the detected cells of the stack (brown dots) (bottom). Blue: DAPI, green: CTB-Af555. Scale bar: 50 µm. (b) Sensitivity and false discovery rate for different thresholds for the nucleus area (Thres. NA) and the averaged prediction scores (Thres. PS). *n* = 3 for each fluorescent dye. Lines mark the mean. (c) Maximum intensity projection (c1) of a stack (CTB-Af555 channel) for one brain slice with the positions of the cells manually detected in the individual layers of the stack (c2, blue) or by the automated cell detection pipeline (c3, green). In c3, positions of cells incorrectly not detected by the automated pipeline (orange) or incorrectly detected by the automated pipeline (red) are marked as well. Scale bar: 100 µm. (d) Proportion of detected cells for two different strategies: automated cell detection pipeline or manual detection on the maximum intensity projection (MIP). *n* = 9 (3 datasets per channel). Lines mark the mean. *p* < 0.001 (***), paired sample t-test. Alt text: Graphical representations of the process and validation of the cell detection pipeline.

### Analysis pipeline for reliable detection of labeled neurons

To determine the positions of labeled thalamic neurons in a large data set consisting of several hundred single images per channel and per animal, we developed a semi-automated cell detection pipeline that uses information from multiple frames of the image stacks to detect cells in the LSCM images (Fig. 2a). In a first step, the positions of the cell nuclei were extracted from the DAPI channel of the LSCM images using StarDist (Schmidt et al., 2018). On average, 93.4 % of all cell nuclei were detected in the test data sets (Supp. Fig. 3). The positions of the cell nuclei were then used to extract image crops that were then classified by previously trained convolutional neural networks (CNNs) as labeled cells or unlabeled cells. The corresponding models had classification accuracies between 96.1 % and 97.4 % for the test data sets (Supp. Fig. 4). Some cells were detected in multiple frames of the stacks and were considered accordingly (Fig. 2a, see Methods). The average prediction scores in each frame and the size of the associated cell nuclei were used as filter criteria. By adjusting the corresponding thresholds, the sensitivity and the false discovery rate (FDR) could be influenced (Fig. 2b). Low thresholds resulted in sensitivities in the range of 76–88%. However, there was still a relatively large number of false positive detections because many more cell nuclei belonged to unlabeled cells (e.g., unlabeled neurons or glial cells). Since high specificity was important, higher thresholds were chosen for the subsequent anatomical studies. These achieved a sensitivity of about 65 % for the single channels. Very few cells were detected false-positive (FDR at about 5 % for the test data sets). If the labeled cells were manually marked on the maximum intensity projection, significantly fewer cells were detected than with the automated cell detection (Mean: 28.2 % vs. 65.1 %, *p* < 0.001, paired sample t-test), indicating that the automated cell detection pipeline is a better strategy compared to a manual annotation of the cells on the maximum intensity projection (Fig. 2c, Fig. 2d).

**Figure 3.**
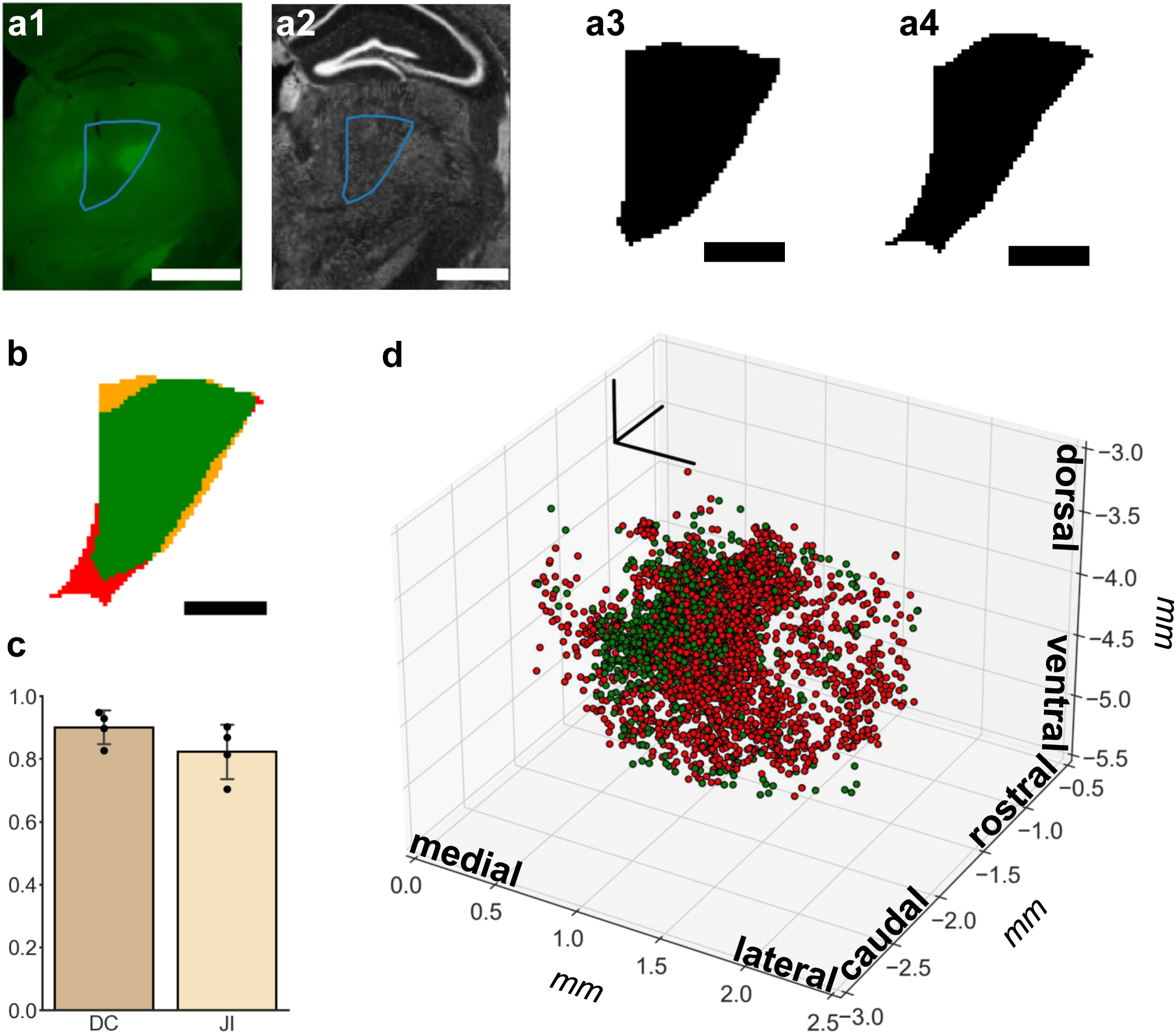
Reconstruction of the cell positions. (a) Manually segmented reference structure. An anatomical structure (here: complex of PO and Eth) was segmented manually on the widefield fluorescence microscope image (blue). The boundaries of the segmentation were then transformed to the CCFv3 (a2) and a binary mask was created (a3). The binary mask of the structure of the corresponding coronal slice of the CCFv3 (a4). Scale bars: 1 mm (a1, a2) and 0.5 mm (a3, a4). (b) Comparison of both binary masks. Pixels are classified as true positive (present in both binary masks, green), false negative (incorrectly missing in the manually segmented and transformed binary mask, red) and false positive (incorrectly present in the manually segmented and transformed binary mask, orange). Scale bar: 0.5 mm. (c) Calculation of the Dice coefficient (DC) and the Jaccard index (JI) for *n* = 4 different manually segmented anatomical structures. Mean ± Standard deviation. (d) Example for a reconstruction of all cell positions detected in the complete data set of the LSCM stacks for one animal (ID32). Position of cells detected after injection of CTB-Af555 in M1 (green) and CTB-Af647 in S1 (red). Alt text: Overview of the process and validation of the reference atlas registration.

**Figure 4.**
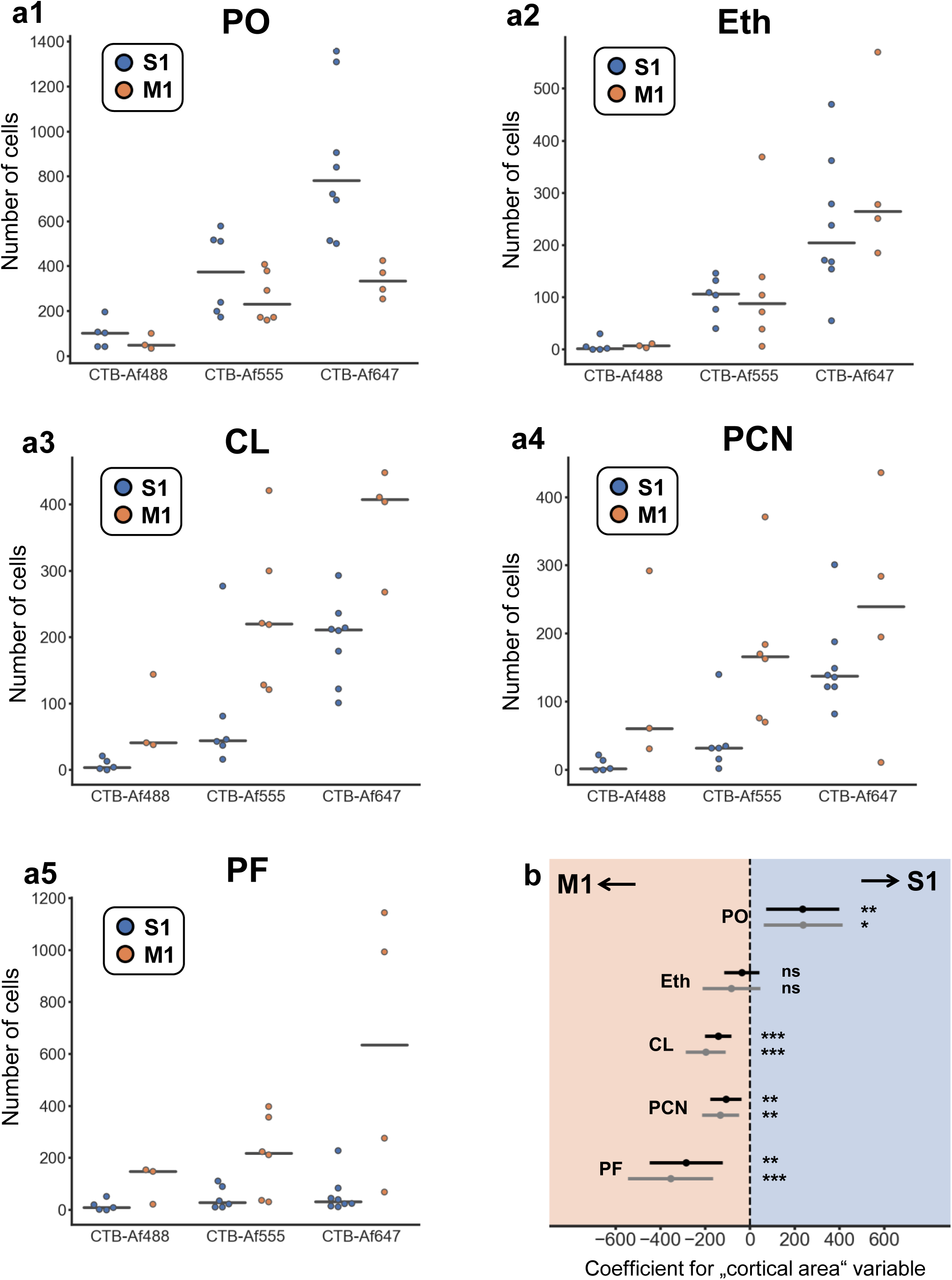
Thalamocortical connection strength to S1 and M1 for different thalamic nuclei. (a) Number of detected cells for *n* = 19 injections in S1 and *n* = 13 injections in M1, stratified by the different fluorescent dyes. Lines mark the median. (b) Coefficients for the *“cortical area”* variable in a multiple linear regression model for each nucleus. Black: number of detected cells, grey: number of detected cells after adjustment for the volume covered by the LSCM stacks. Positive values indicate a stronger projection to S1, negative values indicate a stronger projection to M1. *p* ≥ 0.05 (ns), *p* < 0.05 (*), *p* < 0.01 (**), *p* < 0.001 (***). Alt text: Presentation of the number of detected cells after injections into the primary somatosensory cortex and the primary motor cortex for the analysis of the thalamocortical projection strengths of five thalamic nuclei.

### Reference atlas adaptation ensures high registration accuracy

The positions of the cells were adapted to the CCFv3 reference atlas. This was done by first marking three reference points on the confocal and overview images to transform cell positions from confocal to overview (Supp. Fig. 5a). Secondly, the positions in the overview image were transferred to the reference atlas (Supp. Fig. 5b). This was validated by manual segmentations also employing blood vessel patterns (Fig. 3a, b, Supp. Fig. 6): When the manual segmentation of anatomical structures was compared with the semi-automated adaptation to the CCFv3 (see Methods), performance scores were on average 90 % (Dice coefficient) and 82 % (Jaccard index) (Fig. 3c). The reconstructed cell positions determined in one example mouse are shown in Figure 3d and indicate a medial enrichment of the cell positions in this animal. The data suggest that the atlas registration process resulted in a segmentation with good accuracy.

**Figure 5.**
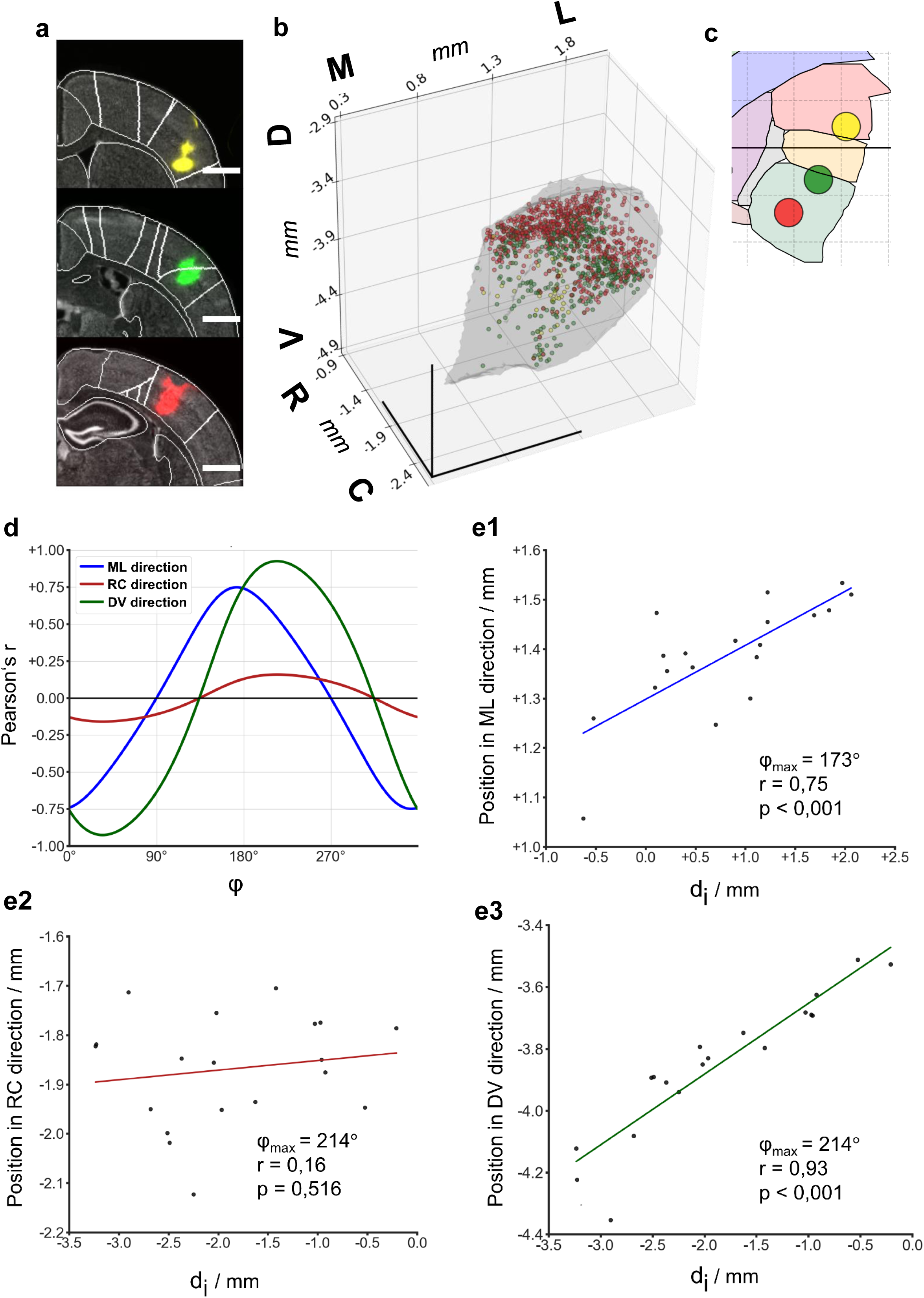
Topography of the cells with respect to the thalamocortical projections from the PO to S1. (a) Injection sites adapted to the corresponding coronal slices of the CCFv3 for one experiment with three injections in S1. Yellow: CTB-Af488, green: CTB-Af555, red: CTB-Af647. Scale bars: 1 mm. (b) Cell positions in the PO. Same colors as in a. M: medial, L: lateral, D: dorsal, V: ventral, R: rostral, C: caudal. (c) Positions of the injection sites (from a) on the two-dimensional representation of the cortex. Same colors as in a. (d) Pearson correlation coefficient for the three different main axis directions for different angles φ.e) Linear regression for the direction (φ_max_) with the highest Pearson correlation coefficient (*r*) for the three different main axis directions. ML: mediolateral, RC: rostrocaudal, DV: dorsoventral. Alt text: Presentation of the analyses of the topographical organization of the neuronal projections from the PO to the primary somatosensory cortex.

**Figure 6.**
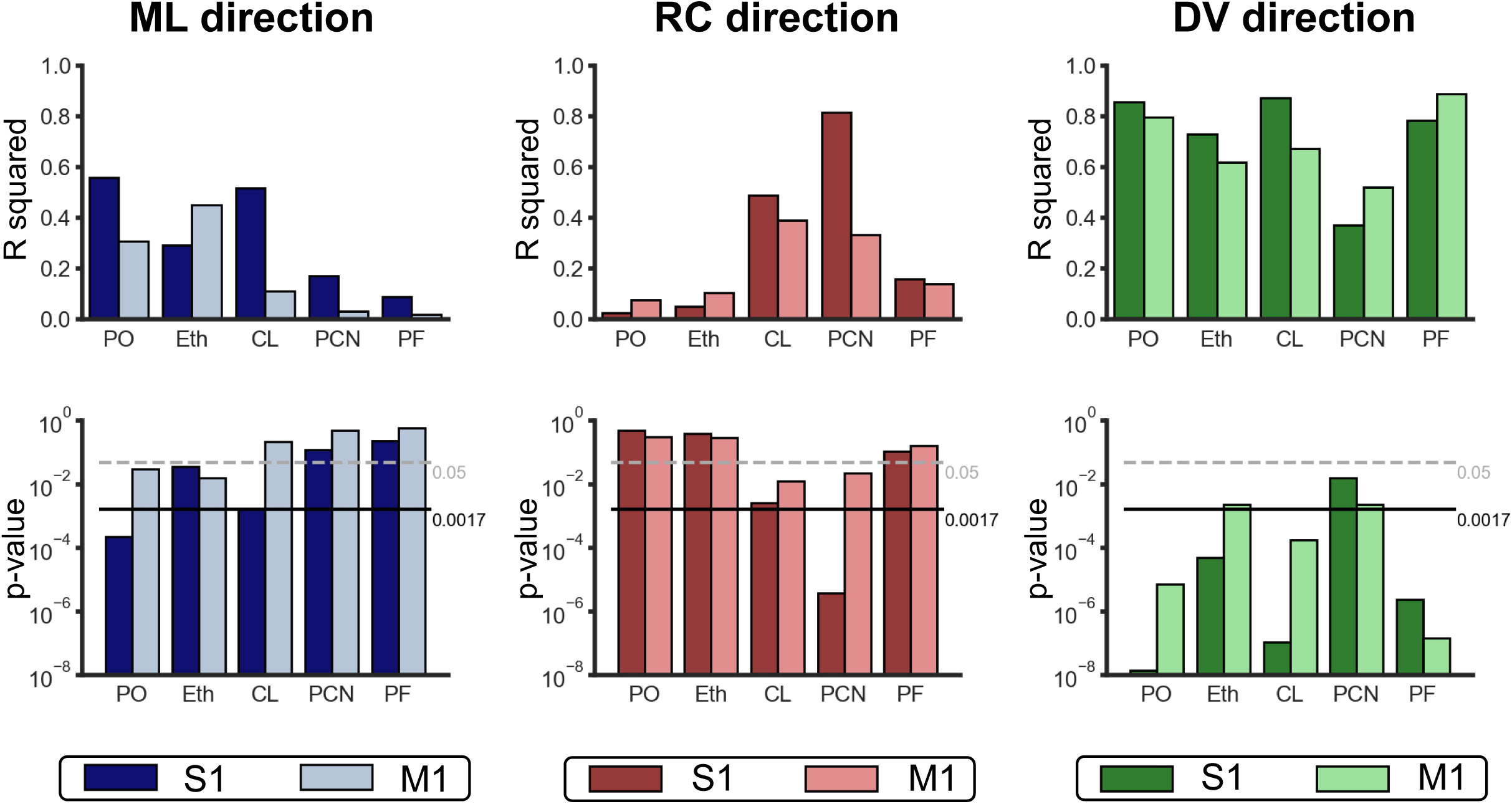
Topographical arrangement of the cells. Topography in the direction φ_max_. Shown are the corresponding *p*-values and the coefficient of determination (*R* squared) for the thalamocortical projections to the primary somatosensory cortex (S1) and the primary motor cortex (M1) for different thalamic nuclei. The values were calculated for the mediolateral (ML), rostrocaudal (RC) and dorsoventral (DV) direction for the five thalamic nuclei. Black continuous line at 0.0017 (Bonferroni corrected significance level), grey dashed line at 0.05 (uncorrected significance level). Alt text: Summary of the topographical analyses of the projections from five thalami nuclei to the primary somatosensory and to the primary motor cortex, presented using bar charts.

Since exact anatomical boundaries of the thalamic nuclei were hardly distinguishable during imaging, the areas microscopically covered were subsequently analyzed in more detail (see Methods). This way the proportion of the volume imaged for each thalamic nucleus could be calculated. Thalamic nuclei were considered in subsequent analyses only if they were covered by at least 50 % in each animal and included PO, CL, PCN, Eth and PF. Furthermore, this was also the case for the ventral posteromedial nucleus (VPM) and the parvocellular part of the ventral complex of the thalamus (VPpc). The VPM was excluded from further studies because labeled axonal bundles within this nucleus made it in parts impossible to clearly identify labeled neurons. The VPpc is divided into a medial and a lateral part in the CCFv3 (Wang et al., 2020). The VPpc contained only very few cells in most of the experiments (median: 4 and 2 cells for injections in S1 and M1, respectively), so it is likely that this area has no relevant thalamocortical connections to S1 or M1. This agrees with previous studies that have described a connection of the VPpc with gustatory and visceral cortical areas (Oh et al., 2014). The VPpc was therefore no longer considered in further analyses. On average, each of the five remaining thalamic nuclei was covered by at least 70 %, and the PO was even covered by almost 100 % in most of the animals (Supp. Fig. 7).

**Figure 7.**
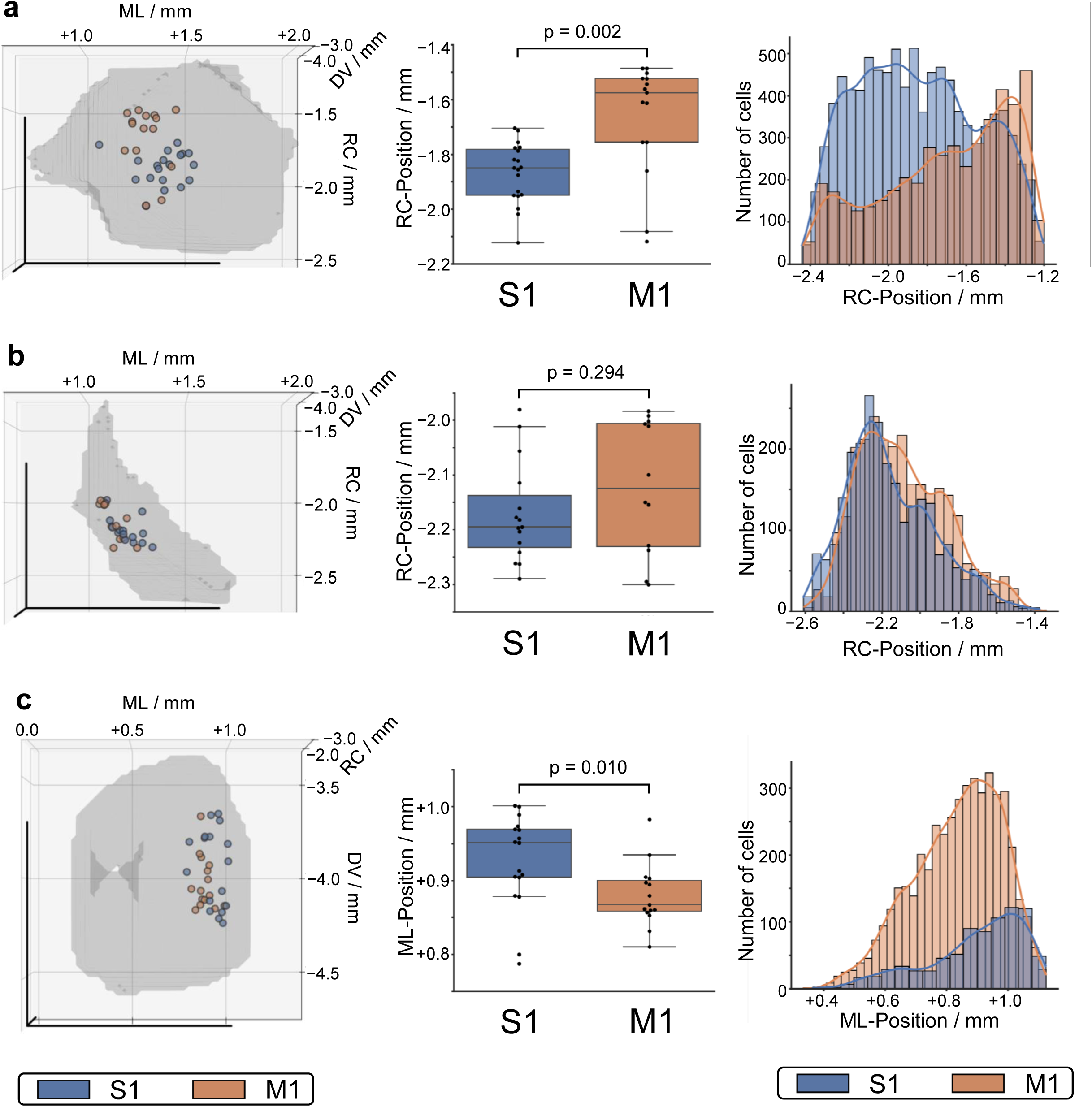
Different distribution of the cells in three thalamic nuclei. (a) PO. (b) Eth. (c) PF. Left: Centroid of the cell positions (each point indicates the centroid of all cell positions for one injection experiment in the primary somatosensory (S1) or the primary motor cortex (M1)). Top view (a and b) and frontal view (c). ML (mediolateral), RC (rostrocaudal), DV (dorsoventral). Middle: Comparison of the position of the centroids in the rostrocaudal (a and b) or mediolateral (c) direction between S1- and M1-injections. Boxplots show the median, interquartile range and whiskers (whiskers indicate all data points within 1.5 times the interquartile range). Right: Histogram for the position of all cells for S1-injections and M1-injections in the rostrocaudal (a and b) or mediolateral (c) direction. Continuous lines are a kernel density estimation for the respective distribution. Alt text: Presentation of the analyses of the different distributions of cells in three different thalamic nuclei detected after injections into the primary somatosensory cortex and injections into the primary motor cortex, using schematic anatomical diagrams, box plots and histograms.

### Structural connectivity to S1 and M1 differs between thalamic nuclei

Injections in S1 or M1 were used to identify the number of neurons in each thalamic nucleus projecting to S1 and M1. Due to differences in the background autofluorescence, different numbers of cells were detected depending on the fluorescent dye used (Supp. Fig. 8). Comparison of the medians stratified for the three different dyes (Fig. 4a) indicated that in the PO more cells were found for S1 injections than for M1 injections. An equal number of Eth neurons projected into S1 and M1, while neurons of the intralaminar nuclei (CL, PCN and PF) targeted the M1 cortex more frequently than the S1 cortex.

**Figure 8.**
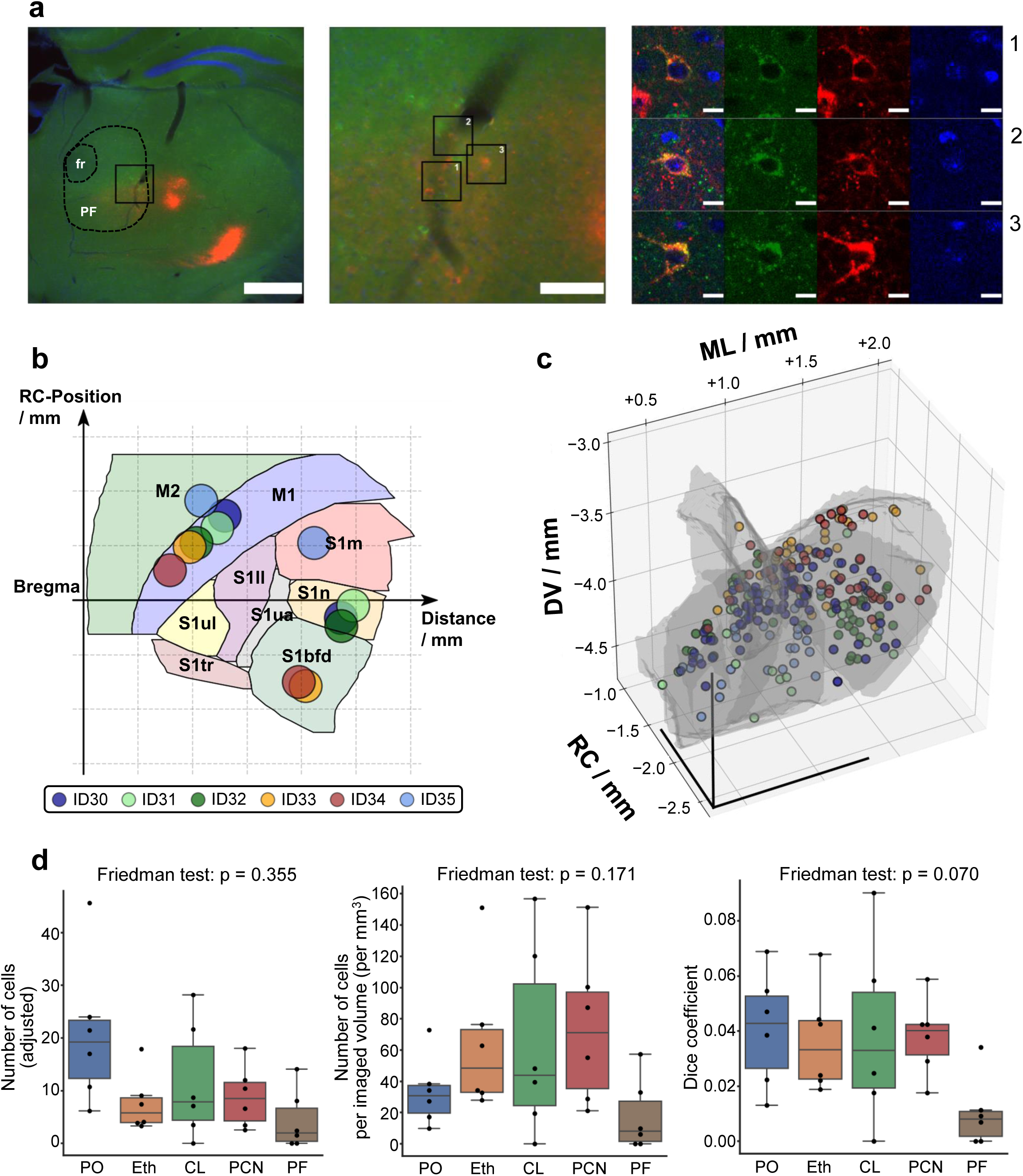
Cells with axonal branching to S1 and M1. (a) Example for cells with uptake of both fluorescent dyes. Left: Overview image (widefield fluorescence microscope image). Middle: Detail from the left image. Right: LSCM images that correspond to the outlined areas from the middle picture. Green: CTB-Af555 channel, red: CTB-Af647 channel, blue: DAPI channel. Dashed lines in the left image show the borders of the PF and the fasciculus retroflexus (fr) (manually segmented). Scale bars: 500 µm (left), 100 µm (middle), 10 µm (right). (b) Positions of the injection sites for the six experiments of the experimental group with co-injections. Each color represents one experiment. Names of the cortical areas as in Fig. 1. RC: rostrocaudal. (c) Cells with uptake of both tracers in the five thalamic nuclei (PO, Eth, CL, PCN and PF) for the six experiments. Colors are the same as in b. DV: dorsoventral, RC: rostrocaudal, ML: mediolateral. (d) Different statistical parameters for the neurons with uptake of both fluorescent dyes. Left: number of cells (corrected for the volume covered by the LSCM stacks), Middle: cell density (number of cells per volume covered by the LSCM images), Right: Dice coefficients. Boxplots show median, interquartile range and whiskers (whiskers are defined in the same way as in Fig. 7) Alt text: Graphical representations of the analysis of the cells with uptake of two tracers used in the co-injection experiments.

To statistically validate these findings, a multiple linear regression model was applied (see Methods). The influence of the chosen injection site (S1 vs M1) on the number of detected cells could be estimated in this way and the corresponding confidence intervals were determined (Fig. 4b). Only injections where 0.2 µl or 0.3 µl of the fluorescent dye has been injected were considered for the analysis of the connection strength, so that the injected volume could be assumed as roughly constant across the experiments. These analyses were also performed for the number of cells after correction for the volume covered by the LSCM stacks. The regression model confirmed that the projection strength to S1 and to M1 differs between the individual thalamic nuclei. The more laterally located PO nucleus projects more strongly to S1 than to M1, the more medially located intralaminar nuclei project more strongly to M1 than to S1. The Eth targets both M1 and S1 to a similar extent.

### Thalamic projection neurons are arranged topographically

To test if neurons within the thalamic nuclei are topographically arranged, we examined their location in the mediolateral, dorsoventral and rostrocaudal direction (hereafter referred to as the main axis directions) within each thalamic nucleus with respect to the position of the injection sites in S1 and M1 (Fig. 5a-c). The direction (indicated as angle φ) that had the strongest positive correlation (i.e., the highest value for the correlation coefficient *r*) between the location of the injection sites and the position of the cells in each of the three main axis directions was designated φ_max_ (Fig. 5d). Therefore, varying the position of the injection site in the direction of φ_max_ had the strongest effect on the position of the cells in the corresponding main axis direction within the respective thalamic nucleus (Fig. 5e). The positions of the injection sites were calculated for different directions (Fig. 6, Supp. Fig. 9, Supp. Tab. 4).

Topographic arrangements of neurons were evident for thalamocortical projections to S1 and to M1 for all five thalamic nuclei. However, the topographical arrangement was different for the different main axis directions and for the different nuclei. For example, the PO showed a clear topographic arrangement with respect to the dorsoventral direction – for projections to S1 as well as to M1 (φ_max_ was 214° (for S1) and 193° (for M1)). Thus, most of the variance of cell positions in dorsoventral direction in the PO can be attributed to the variance of the position of the injection site in the corresponding direction (S1: *r^2^* = 0.86, *p* = 1.39 x 10^-8^; M1: *r^2^* = 0.80, *p* = 7.21 x 10^-6^). It can be concluded that more dorsally located areas within the PO project to more caudal subareas within S1 and within M1 because φ = 180° is defined as the direction pointing in caudal direction (see Supp. Fig. 9) and both values for φ_max_ are close to this value. In contrast, the positions of the injection sites within S1 or M1 seem to have no or only a minor influence on the rostrocaudal position of the cells in the PO (S1: *r^2^* = 0.03, *p* = 0.516; M1: *r^2^* = 0.08, *p* = 0.315). A distinct topographical arrangement of cells in the dorsoventral direction was also evident for the PF (S1: *r^2^* = 0.78, *p* = 2.35 x 10^-6^; M: *r^2^* = 0.89, *p* = 1.44 x 10^-7^). However, the choice of the injection site for the PF seems to have no or only little effect on the position of the cells in the mediolateral direction for the PF (S1: *r^2^* = 0.09, *p* = 0.243; M: *r^2^* = 0.02, *p* = 0.603). Thus, the positions of the neurons within the thalamic nuclei indicate a topographic arrangement with respect to the thalamocortical projections.

### Differential distribution of cells projecting to both the primary somatosensory cortex and the primary motor cortex

Consequently, in certain directions, the position of the injection sites within S1 or within M1 has little or no effect on the positions of the cells regarding these directions. The PO and the Eth – for the rostrocaudal direction – and the PF – for the mediolateral direction – had low coefficients of determination (*r^2^* < 0.15) for S1 as well as for M1. We tested whether the choice of the cortical area, i.e., whether the injection was previously made in S1 or in M1, influenced the positions of the cells for these three nuclei. In fact, there was a significant difference for the PO (rostrocaudal direction, *p* = 0.002) and the PF (mediolateral direction, *p* = 0.010) (Fig. 7). For the Eth, the corresponding difference was not significant (rostrocaudal direction, *p* = 0.294). Thus, in the PO, cells projecting to M1 tend to lie more rostrally than cells projecting to S1. In the PF, cells projecting to M1 lie more medially than cells projecting to S1. The choice of the injection site within S1 or within M1 had no large influence on the position of the cells as it is indicated by the low (and not-significant) coefficients of determination. Thus, the different positions of the injection sites within S1 and within M1 cannot fully explain this difference. Consequently, neurons projecting to S1 appear to be distributed differently within the corresponding thalamic nuclei compared to neurons projecting to M1.

### Cells with simultaneous projections to the primary somatosensory and the primary motor cortex

To test if thalamic neurons project to both S1 and M1 cortices, co-injections of CTB with different colors were placed in both cortical areas (Fig. 8a, Supp. Fig. 10). The topographical arrangement of the thalamocortical projections identified above was used to define injection sites in a way to maximize the chance of finding cells with uptake of both fluorescent dyes (Fig. 8b). The two tracers CTB-Af555 and CTB-Af647 were used because a large number of neurons could be detected (see Fig. 4, Supp. Fig. 8). Neurons with uptake of both fluorescent dyes were found in all six animals and in all five thalamic nuclei (Fig. 8c,d). However, the number of cells that had taken up both fluorescent dyes was small compared to the number of cells that had taken up only one fluorescent dye. This is also reflected in the low Dice coefficients (Fig. 8d, right). The Friedman test, which was used to determine differences between the groups, was not significant for the variables examined (p > 0.05) (Fig. 8d).These results indicate that neurons with axonal branching and axonal terminals in S1 and M1 are a general principle throughout the higher-order thalamus.

## Discussion

The connections of several higher-order thalamic nuclei to two different cortical areas – the primary somatosensory and primary motor cortex – were investigated using a cell detection pipeline based on CNNs. About twice as many PO neurons were connected with the S1 compared with the M1 cortex in our experiments. Projection neurons within several thalamic nuclei were topographically organized with regard to their projection targets. Moreover, co-injections revealed neurons with a simultaneous projection to S1 and M1 in all five thalamic nuclei investigated. Hence, our work provides important anatomical insights into the structural and organizational principles of connections between higher-order thalamic nuclei and the cortex. These findings may be applied to other thalamic nuclei or cortical areas and may thus represent a generalizable design principle of thalamocortical projections.

To achieve a quantitative description of the connection patterns of the PO and surrounding nuclei with S1 and M1, a volume of more than 2 mm^3^ was imaged per animal with a voxel size of 0.38 µm x 0.38 µm x 2.5 µm in the region of the right thalamus. Analysis of this vast amount of data required an automated cell detection algorithm. Detection of cells using this 3D approach turned out much more efficient than using a manual detection approach based on maximum intensity projections of the stacks. This indicates that important information might get lost when using a maximum intensity projection of a stack instead of the individual frames of a stack. The cell detection pipeline also considered the possible detection of cells in multiple frames and by adjusting appropriate threshold values, sensitivity and false discovery rate could be improved. The cell detection pipeline can also be used for the detection of cells to address other research questions or can – by training the CNNs accordingly – be used for the detection of other structures.

The analysis of the data showed that the thalamic nuclei differ in their structural connectivity to S1 and M1. This could be related to functional differences between the thalamic nuclei, such as the PO to S1 are stronger than to M1, simply because PO has a role in the somatosensory system (Mease & Gonzalez, 2021; Van Horn & Sherman, 2007). However, it must also be emphasized that only a small area of the respective cortical area was covered by each injection. This may also explain why fewer cells were found per volume after injections in S1 than after injections in M1.

Moreover, thalamic nuclei do not project uniformly to the different layers of the cortex. The PO, for example, projects primarily to layer 1 and layer 5a in S1 (Jouhanneau et al., 2014; Mease & Gonzalez, 2021). Therefore, the exact distribution of the fluorescent dye in the cortex also affects the number of cells detected. And thalamic neurons may not project to all subareas of S1 or of M1 equally. Furthermore, CTB has been reported to be taken up directly via the axolemma and not only via synaptic terminals (Chen & Aston-Jones, 1995), although our own results do not support this (Körber et al., 2014). Uptake along the axon would bias the location-specificity of the injection site, because also neurons whose axons traverse the injection site without forming synapses would be labeled.

The comparison of the different injections in S1 and different injections in M1 showed that the projection neurons are highly topographically arranged. Topographical arrangements are a general construction principle of the nervous system, which is also established across animal species: for example, the motor cortex of mammals is structured with a relation to different areas of the body (Levine et al., 2012), or single barrels in the rodent barrel cortex can be assigned to individual whiskers (Alloway, 2007). In rats, it has already been shown that neurons in the PO or POm show a topographical arrangement with respect to projections to S1 and S2 (Fabri & Burton, 1991). Different subareas of the thalamic nuclei could thus possibly have different functional properties. However, a topographical arrangement also makes sense for reasons of efficiency: it leads to axonal connections that are shorter, which saves space and energy (Chklovskii & Koulakov, 2004; Patel et al., 2014). This could also explain why the more laterally located PO has stronger thalamocortical connections with the more laterally located S1 and the more medially located intralaminar nuclei have stronger thalamocortical connections with the more medially located motor cortex This organization reduces the total length of axonal connections. A characterization of the topographical arrangement as performed in this study may also help to plan further experiments. For example, researchers could use these findings to label cells in desired subareas of the thalamic nuclei.

Neurons projecting to S1 and neurons projecting to M1 are not distributed equally in the thalamic nuclei. Cells projecting to S1 tend to be located more laterally than cells projecting to M1 in the PF. In the PO, cells projecting to M1 are located more rostrally than cells projecting to S1. The uneven distribution could be related to different substructures within the thalamic nuclei: the murine PF can be divided into a medial, a central and a lateral part (Mandelbaum et al., 2019). This division is based on findings about differences in transcriptional, electrophysiological, and anatomical properties (Mandelbaum et al., 2019). Sumser et al. (2017) divided the posterior complex of the thalamus in four subnuclei because anterograde tracing experiments showed segregated clusters of giant boutons. They named these subnuclei anterior PO (POa), POm_medial_, POm_lateral_ and PoT (triangular part of the PO) (Sumser et al., 2017). The PoT is segmented as an independent thalamic nucleus in the CCFv3 (Wang et al., 2020). The Pom_medial_ is possibly identical to the Eth in the CCFv3. In this case, the PO – as it is defined in the CCFv3 – consists of a more rostral part (POa) and a more caudal part (defined as POm_lateral_ by Sumser et al.). Hence, our work suggests that cells projecting to the motor cortex lie in more rostral areas and therefore predominantly in the POa. Other studies indicate that the POm is divided in substructures along the rostrocaudal axis (El-Boustani et al., 2020; Ohno et al., 2012). This may align with the division in the POa and the POm_lateral_, although the latter may be renamed as posterior part of the PO (POp). However, further research is required to obtain a more precise anatomical and functional differentiation of individual subnuclei of the posterior complex of the thalamus.

The co-injections revealed that all five higher-order thalamic nuclei contain neurons with a simultaneous projection to S1 and to M1. The cortical injection sites were chosen based on our delineation of the topographical arrangement to maximize chances to obtain dually labeled projection neurons. Nevertheless, the number of cells that have taken up both fluorescent dyes was low compared to the number of cells labeled by only one of the fluorescent dyes. In other experiments with co-injection of retrograde tracers into S1 and S2 in rats, only a few cells with uptake of both tracers were detected as well (Spreafico et al., 1987). This can also be attributed to the fact that only a small area of S1 or M1 can be covered by the injection sites. Finally, it cannot be inferred from our method how many cells have simultaneous projections to S1 and M1. Not all neurons will have bifurcated axons targeting both cortical areas. While cells projecting to S1 and M1 are not evenly distributed in the thalamic nuclei, single cell tracing analyses in rats showed that in the POm, some, but not all cells examined, had axonal terminals in both the primary somatosensory cortex and the motor cortex (Ohno et al., 2012). However, all cells reconstructed had terminals in S1 and in other cortical areas (Ohno et al., 2012). Cells with axonal terminals in multiple cortical areas thus seem to be a general building principle within the thalamus and are probably linked to a functional role. Such cells could be involved in the synchronization of different cortical areas (Saalmann, 2014; Saalmann et al., 2012). Branching of axons with terminals in more than one cortical area would have the potential to transmit the same information to several cortical areas (Sherman, 2016).

### Technical limitations of the study

The training and validation of the cell detection was based on a manual classification of the cells. It should be noted here that, for some of the cells, it was difficult to clearly identify whether the cell was labeled or not. This could be due to a weak fluorescence signal, or to the fact that, in some cases, it was not possible to clearly distinguish between labeling of a cell and labeling of axons passing close to the cell nucleus.

During injections into the motor cortex, most injections were made in the medial and, with respect to the rostrocaudal direction, middle regions of M1. This corresponds most closely to the areas of M1 associated with the whiskers (M1wk) (Casas-Torremocha et al., 2022; Vanni et al., 2017). However, this subregion was not delineated in the CCFv3. That most of the injection sites were located medially in M1 had the advantage of reducing the likelihood that a significant amount of the fluorescent dye could diffuse into the adjacent S1. Nevertheless, M1 is less completely covered by injection sites than S1, which needs to be considered when interpreting the data. It must also be noted that diffusion of the dye into the adjacent secondary motor cortex (M2) cannot be ruled out. For three injection sites, based on the segmentation using the CCFv3, it can also be assumed that a larger proportion of the dye is located in M2. This should be taken into account as a limitation when interpreting the results.

The method of the adaptation to the reference atlas has also methodological limitations. A wrong assignment of detected cells is – especially in the peripheral area of the thalamic nuclei and in particular for smaller nuclei – possible. Coronal reference slices of the CCFv3 were selected for each brain slice. However, the positioning of the brain in the microtome inevitably resulted in tilting, so that no brain slice corresponded precisely with a coronal slice of the reference atlas. To minimize the resulting effects, landmarks especially of the right-sided thalamus were used for the selection of the appropriate coronal reference atlas slice since this was the area of interest for the anatomical studies. Nevertheless, incorrect assignment of cells cannot be ruled out. For example, tilting could have resulted in incorrect assignment of cells located in the mediodorsal nucleus (MD) to the PF and vice versa. Therefore, peripheral areas of the thalamic nuclei or smaller nuclei have a higher risk of incorrect assignment of detected cells. The PO itself as a rather large thalamic nucleus is probably less affected by incorrect assignment.

It should also be pointed out again that only the PO was almost completely covered by the LSCM stacks, while the other thalamic nuclei were covered >50 % in each data set. In particular, there was an underrepresentation in the areas at the edges of the microscopic images (e.g., the medial region of the PF or caudal regions of the Eth). When examining the connection strength of the individual nuclei to S1 and to M1, similar values were found for values that have been corrected for the proportion of the volume covered by the LSCM stacks and values that have not been corrected. Consequently, it can be assumed that incomplete coverage of the thalamic nuclei probably had only a minor influence on the results.

This study provides important foundations for the organization principles of thalamocortical projections to S1 and M1. The insights gained could potentially be transferable to other thalamic nuclei and other cortical areas, thus representing a general building principle for cortico-thalamocortical networks.

## Methods

### Animals

Experiments have been approved by the Regierungspräsidium Karlsruhe (Badenwuerttemberg, Germany) under the reference number 35-9185.81/G194/15. Male mice (C57/BL6 stain) were between 11 and 25 weeks old and bought form Charles River Laboratories. Efforts were made to minimize suffering of the animals. Mice were kept under controlled environmental conditions (i.e., controlled temperature, humidity, and light-dark cycle) and food and water were available ad libitum.

### Injection of the fluorescent dyes

All injections were made in the right cerebral cortex (S1 or M1) (Fig. 1b). Stereotactic coordinates were determined with the help of a reference atlas (Franklin & Paxinos, 2001). Mice were anesthetized by intraperitoneal injection of an anesthetic mixture (injected volume: 3 µl per gram body weight plus additional 10 µl). The mixture consisted of Medetomidine (1 mg/ml), Fentanyl (0.05 mg/ml) and Midazolam (5 mg/ml), which had been mixed in a volume ratio of 3 : 2 : 8. After shaving of the scalp, the skin was disinfected and local anesthesia was applied (subcutaneous and epicutaneous application of lidocaine hydrochloride, 1 %). An ophthalmic ointment was applied to prevent drying out. As soon as a sufficient depth of anesthesia was reached (additional 40 µl of the anesthetic mixture were injected if necessary) and no reactions to pain stimuli were detectable, the mouse was positioned in a stereotactic apparatus. After preparation of the skull surface, the skull was leveled. Bregma, lambda and two points (1 mm to the right and to the left of bregma) served as orientation points for levelling. For some of the injections (S1 injections of the experimental group with co-injections), the skull was rotated 20° to the left side before the injection to allow injection as perpendicular to the cortex surface as possible (similar to a previous report (Sumser et al., 2017)). After rotation of 20° the position of the skull was adjusted so that bregma and lambda had again the same height and both were inside the sagittal plane. The skull was drilled over the planned injection sites. Glass capillaries, pulled with a micropipette puller, were filled with the respective fluorescent dye (CTB-Af488, CTB-Af555 or CTB-Af647) and positioned at the coordinates within the cortex that have previously been determined. Therefore, 100 µg of the fluorescent dye (Invitrogen) was dissolved in 100 µl phosphate-buffered saline (PBS). Bregma served as a reference for determining the mediolateral and rostrocaudal position, whereas the cortex / dura surface served as a reference for determining the dorsoventral position. After a short waiting period, 0.2 to 0.5 µl of the fluorescent dye was slowly injected. After a waiting period of at least two minutes, the glass capillary was removed from the brain. After the injections, the skin incision was sutured and 300 µl Carprofen (0.5 mg/ml) was given subcutaneously. The anesthesia was then antagonized with Atipamezole (5 mg/ml), Flumazenil (0.1 mg/ml) and Naloxone (0.4 mg/ml) (volume ratio: 3 : 40 : 6, 150 µl intraperitoneally and 100 µl subcutaneously).

### Tissue preparation

Mice were deeply anesthetized by intraperitoneal injection of Pentobarbital (3.125 µl/g body weight, additional 50 µl if necessary). After ensuring a sufficient depth of the anesthesia, the mouse was perfused transcardially with PBS followed by paraformaldehyde solution (PFA, 4 %). The brain was extracted and postfixed for approximately 24 h in PFA (4 %) in a fridge and rinsed several times with PBS and subsequently cut with a microtome into 80 µm thick slices. The slices were transferred to a PBS solution that contained 4’,6-Diamidino-2-phenylidol (DAPI) for cell nuclei staining and then mounted on slides (mounting medium: polyvinyl alcohol).

### Widefield fluorescence microcopy

Overview images were taken with a wide-field fluorescence microscope (Leica DM6000). The PO should be covered as completely as possible by the microscopic images. About 17 consecutive brain sections were recorded accordingly. The signal was acquired with a 10x objective, which had a numerical aperture of 0.40. Multicolor LED illumination, filters, exposure time and intensity levels were selected appropriately for each channel. Tile merging was done by the software of the microscope.

Additionally, overview images of the brain slices that contain the signal from the injection site were taken (1.25x objective, numerical aperture: 0.04), including an additional bright field image. The image, on which the injection site was most clearly visible, was used for the subsequent adaptation to the reference atlas to determine the exact location of the injection site. The images of the injection site were also used to evaluate the quality of the injection: Injection experiments were excluded from the anatomical analyses if the injection was too deep, if there was a strong reflux of the dye onto the brain surface or if the signal intensity was low. Animals were also excluded if data from the consecutive brain slices could not be obtained completely, for example, due to tissue damage during the slicing process. In total, *n* = 19 injections in S1 and *n* = 15 injections in M1 could be used for the anatomical studies. However, injections excluded from the anatomical studies may have been used to train and test the CNNs for the cell detection pipeline.

### Laser scanning confocal microscopy

Detailed image stacks (*z*-spacing: 2.5 µm) of the area of interest were captured by laser scanning confocal microcopy (LSCM, Leica SP8). For this purpose, an overview image of the right hemisphere of each section was taken at low resolution and the region of interest, which should include the PO as complete as possible, was defined. The region of interest was defined by comparison of anatomical landmarks with two reference atlases (Dong, 2008; Franklin & Paxinos, 2001). An exact delineation of the borders of the thalamic nuclei was not possible on the overview images. The volume covered by the LSCM stacks was therefore calculated afterwards (see below). Different laser wavelengths were used for the different signals (DAPI: 405 nm, CTB-Af488: 488 nm, CTB-Af555: 552 nm, CTB-Af647: 638 nm). A 40x lens (numerical aperture: 1.1, immersion medium: water), an additional zoom factor of 0.75x (set in the software) and a resolution of 1024 x 1024 pixels resulted in a pixel size of 0.38 µm x 0.38 µm. For the training of the StarDist neural network, images were also used that had been taken with slightly different settings (20x lens, immersion medium: oil, additional zoom factor: 1.5x). However, these settings resulted in the same pixel size.

The DAPI and CTB-Af488 signals were detected with photomultiplier tubes and the CTB-Af555 and CTB-Af647 signals with hybrid detectors. In each case, suitable wavelength ranges were selected for the detection of the emitted photons and left constant throughout the experiments. The laser intensity was selected appropriately. To reduce the strong autofluorescence signal in the CTB-Af488 channel, an offset of -7.0 % was set for this channel. The scan was conducted as a bidirectional and sequential scan. The DAPI signal and the CTB-Af647 signal could be recorded during the same scan. Tile merging was again performed by the software of the microscope. Fiji (Schindelin et al., 2012) was used to check the LSCM stacks and remove upper or lower layers of the stacks that contained no or little signal of the brain slices.

Possible crosstalk between the channels should be excluded (Supp. Fig. 1). For this purpose, each of the dyes was at first injected in S1 of one animal and one brain slice was imaged and analyzed with the settings used for the LSCM. Labeled cells were segmented in the corresponding channel. An adjacent area – without signal from the fluorescent dye – was also marked to calculate a relative signal intensity from the ratio of the average signal intensities of both areas. The relative signal intensities were also analyzed for the other two channels for the same areas in order to exclude crosstalk.

### Cell detection

The pipeline developed for cell detection consisted of several steps (Fig. 2a). In the first step, cell nuclei were detected using StarDist (Schmidt et al., 2018) (Supp. Fig. 3). A StarDist network was trained with training data generated manually by using Labkit (Arzt et al., 2022). To evaluate the performance of the detection of the cell nuclei, the automated StarDist cell detection was compared with a manual registration for several image crops (four images, each 800 x 800 pixels).

Only a small proportion of the cells detected by StarDist belonged to labeled neurons. Therefore, the position of the cell nucleus was used as a seed and the surrounding 60 x 60, 90 x 90 or 120 x 120 pixel field could be extracted as an image crop. The extracted image crops were then classified as “cell” (i.e., neuron that had taken up the fluorescent dye) or “no cell” by using Convolutional Neural Networks (CNNs) that were specific to the respective size of the extracted image crops and the corresponding channel. The CNNs were trained by making manual marks for “cells” and for “no cells” (Supp. Fig. 4a) for each of the three (CTB-Af488, CTB-Af555, CTB-Af647) channels in Fiji (28 crops of different stacks for each channel). Subsequently, image crops with the above three sizes were extracted. Care was given to mark image areas that were potentially difficult for the CNNs to distinguish. The images were also rotated (90°, 180° or 270°) and / or mirrored to generate more training data. Finally, some images were randomly removed from the training data set so that the same number of extracted images were classified as “cell” and as “no cell”. The image data was transformed (rescaling – 60 x 60 pixels to 30 x 30 pixels, 90 x 90 pixels to 45 x 45 pixels and 120 x 120 pixels to 40 x 40 pixels – and normalized so that the pixel values had a mean of 0 and a standard deviation of 1). The CNN itself was a binary classifier with two pooling layers and one fully connected layer. An additional dropout (dropout rate: 0.5) should prevent overfitting (Hinton et al., 2012). Binary cross entropy was used as loss function and the activation function of the output layer was a sigmoid function. Nine different CNNs (three different channels and three different formats) were trained (number of epochs: 30, learning rate: 5·10^- 5^). To validate classification accuracy of the CNNs, test data were obtained in a similar way as the training data. For this purpose, crops from seven brain slices, which had not been used for the training of the CNNs, were used.

In order to minimize false positive detections (i.e., classification as a “cell” even though “no cell” would be the correct label), the following strategy was used: first, for each via StarDist detected cell nucleus, a 60 x 60 pixel large image crop was extracted. All those extracted images were excluded that had relatively low signal intensities (first: 95th percentile pixel value of the extracted image was lower than the median of all 95th percentiles for all extracted images of the corresponding layer, or second: mean signal intensity of the extracted image was lower than 800 – for the CTB-Af488 channel and the CTB-Af555 channel – or 100 – for the CTB-Af647 channel). For the not excluded images, a prediction was then made based on the corresponding CNN. For images with a prediction score over 0.3, the 90 x 90 pixel field and 120 x 120 pixel field were extracted as well and classified. Then an average value was calculated from all three prediction scores (average prediction score, APS). Images were not considered further if the APS was below 0.5. The rest of the images were further considered.

A few cell nuclei were detected multiple times in the same layer of the stack (due to oversegmentation by the StarDist neural network). Therefore, if two seeds in a layer had a distance of less than 8 µm, only the one with the higher APS was considered further. Furthermore, due to the rather low *z*-spacing of 2.5 µm, some cells were detected in more than one layer. Therefore, seeds belonging to the same cell were combined by assuming that seeds in different layers with a distance of less than 12 µm belong to the same cell. In this way, several APS values could be assigned to a potential cell. Potential cells were excluded if the area of their nucleus (determined by the segmentation results of StarDist) was smaller than 400 pixels. A potential cell (characterized by a set of APS values) was only classified as such if it had at least one, at least two or at least three APS values that were greater than 0.95, 0.75, 0.50, respectively.

A validation of the final cell detection pipeline was done by manually labelling cells in several crops of stacks (three stacks per channel; the data were from animals not used for the training or testing of the CNNs). Here, the images were selected so that they contained a relatively large number of labeled neurons and so that these labeled neurons could be classified as “cell” or as “no cell” as unambiguous as possible. The distances of the cells labelled in this way were then compared in an automated way by interpreting points with less than 10 µm distance between the manual and the automatic annotation as the same cell. By comparison with the manual annotation, the cells detected by the cell detection pipeline could then be classified as true positive, false positive, or false negative detections.

Cells with uptake of both fluorescent dyes were determined by analyzing which cell positions of the cells detected in the CTB-Af555 channel and the CTB-Af647 channel had a distance below 10 µm. The cells identified in this way were then screened manually and those that did not correspond to a cell with uptake of both fluorescent dyes (e.g., due to false positive detection) were excluded.

### Adaption to the reference atlas

The data sets for the voxel size 25 µm x 25 µm x 25 µm for the CCFv3 were downloaded from the website of the Allen Brain Institute. The position of bregma was defined so that bregma matched the definition of bregma in the Paxinos and Franklin atlas (5.70 mm right, 0.10 mm dorsal and 5.35 mm caudal from the upper, rostral, left corner of the CCFv3).

The two-dimensional representation of the cerebral cortex was created by determining the distance of the boundaries of the cortical areas to the longitudinal cerebral fissure for every fourth coronal slice of the CCFv3 (Supp. Fig. 2a and Supp. Fig. 2b). This distance was used to define the mediolateral position for the two-dimensional representation of the cortex. The rostrocaudal position resulted from the position of the corresponding coronal slice. In this way, the boundaries of the cortical areas were definable for the two-dimensional representation of the cortex.

To determine the position of the individual injection sites, the brain slice which showed the signal of the injection site most clearly was selected. The brightfield image was then used to select an appropriate coronal slice of the CCFv3 – based on anatomical landmarks. Corresponding pairs of points on the microscopic image and the reference atlas slice were then used to transform the microscopic image to the reference atlas by using the Fiji plugin BigWarp (Bogovic et al., 2016) (transformation by the “thin plate spline” method) (Supp. Fig. 2c, Supp. Fig. 2d, Supp. Fig. 2e and Supp. Fig. 2f). The position of the injection site in relation to the two-dimensional representation of the cortex was then determined from the position of the selected reference atlas slice (rostrocaudal position) and from the distance of the injection site from the longitudinal cerebral fissure (mediolateral position) using Fiji.

The adaptation of the cell positions to the reference atlas was performed in two steps (Supp. Fig. 5):

In a first step, three corresponding reference points were defined on the average intensity projection of the LSCM images (CTB-Af555 channel) and on the overview images taken by widefield fluorescence microscopy (CTB-Af555 channel). These points were determined based on easily identifiable structures in both images (e.g., points at branches of blood vessels). The coordinates of the position of a detected cell in relation to the LSCM images could in this way be transformed to the overview images:

Let (*l_0_, o_0_*), (*l_1_, o_1_*) and (*l_2_, o_2_*) be the pairs of reference points for the LSCM images (*l_i_*) and the overview images (*o_i_*) (the three points of both images do not lie on the same line). For the LSCM image, let *L* be a vector space over ℝ*^2^* with *l_0_* as zero vector and for the overview image, let *O* be a vector space over ℝ*^2^* with *o_0_* as zero vector. A detected cell on the LSCM image is now defined by *p_L_ =* λ*_1_l_1_ +* λ*_2_l_2_*(*Eqn. 1*) and we search for the corresponding vector *p_O_* on the overview images. We can assume that *f: L* → *O* is an affine transformation and therefore *p_O_ =* λ*_1_o_1_ +* λ*_2_o_2_* (*Eqn. 2*). Finally, *p_O_*can be calculated since λ*_1_* and λ*_2_* can be derived from *Eqn. 1* as solution of the corresponding system of linear equations.

The validity of this approach was demonstrated qualitatively by manually delineating blood vessels on the average intensity projection of the LSCM stack for three different brain slices and then transforming them in the described way (Supp. Fig. 6).

In the second step, a corresponding coronal atlas slice of the CCFv3 was selected for each overview image by using anatomical landmarks. Then, the right diencephalon was marked on both images (the overview image – here again the CTB-Af555 channel was used – and the reference atlas slice) with matching pairs of points. The ImageJ plugin BigWarp (Bogovic et al., 2016) was then used to transform the points from the overview images to the images of the CCFv3 by using the “thin plate spline” transformation method.

To evaluate the accuracy of the adaptation to the reference atlas, anatomical structures that could visually be well delineated (either VPM or the complex of PO and Eth) were segmented manually and the resulting boundary points were transformed to the reference atlas in the described way (Fig. 3a). By comparing the segmentation of the structures in the CCFv3 with the manual segmentation (after the application of the transformation), the quality of the adaptation was assessable (Fig. 3b).

The assignment of the detected cells to the individual anatomical regions of the CCFv3 was done by defining a list of all regions the voxel belonged to. The region a detected cell could be assigned to then resulted from the voxel to which the cell belonged to.

Since the anatomical regions could not be clearly delineated during the acquisition of the LSCM images, the exact area covered by the LSCM stacks was calculated in another step after the imaging (Supp. Fig. 7). For this purpose, the boundaries for the average intensity projections of the LSCM stacks (CTB-Af555 channel) were defined manually. The boundaries were then transformed to the overview images (widefield fluorescence microscopy images) and to the CCFv3 according to the above algorithm. The so transformed area of the CCFv3 was then converted into a binary mask. The binary mask was then compared with the binary mask of the structure of the selected coronal slice of the CCFv3. By comparing all LSCM average intensity projections with the corresponding CCFv3 segmentations, the volume covered by the LSCM stacks was estimable for each thalamic nucleus and each animal.

### Software

For the image processing and analysis Fiji (Version 1.53c) (Schindelin et al., 2012) and the Fiji-Plugins Labkit (Arzt et al., 2022), StarDist (Schmidt et al., 2018) and BigWarp (Bogovic et al., 2016) were useful. A script was downloaded from the Website of BigWarp and adapted for the purposes in this study. Python (Version 3.7) (Van Rossum, 1995), the Python standard library and Tensorflow (Abadi et al., 2015), Keras (Chollet, 2015), Pandas (McKinney, 2010), Numpy (Harris et al., 2020), Requests (Reitz, 2023), Pynrrd (Everts, 2022), Pillow (Umesh, 2012), Scikit-Image (Van der Walt et al., 2014), Read-roi (Mary, 2020), Shapely (Gillies et al., 2007), Scikit-Learn (Pedregosa et al., 2011), Scipy (Virtanen et al., 2020), Statsmodels (Seabold & Perktold, 2010), Matplotlib (Hunter, 2007) and Seaborn (Waskom, 2021) were used for analysis, statistical tests and presentation of the data. Analysis data were saved as “.xlsx” (Microsoft Excel spreadsheets), “.csv” or “.txt” files. Neural machine translation (https://www.deepl.com/) was helpful in writing and formulating the text of the manuscript.

### Statistics

Dice coefficient and Jaccard index to evaluate the performance of the adaption to the reference atlas were calculated by:

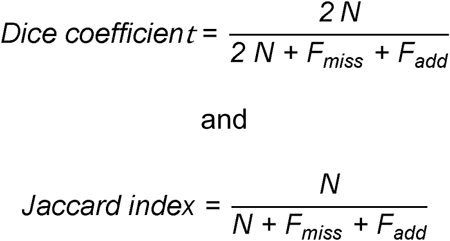

(*N*: number of pixels present in both binary masks, *F_miss_*: number of pixels missing in the binary mask of the manual annotation (compared to the binary mask of the CCFv3), *F_add_*: number of pixels that are additionally present in the binary mask of the manual annotation (compared to the binary mask of the CCFv3)).

To calculate the connection strength of different thalamic nuclei to S1 and to M1, a multiple linear regression model was used. The number of detected cells in the respective thalamic nucleus (or the number of detected cells adjusted for the volume of the thalamic nucleus covered by the LSCM stacks) was predicted in the model by the two independent variables *“used tracer”* (CTB-Af488 vs. CTB-Af555 vs. CTB-Af647) and *“cortical area”* (cortical area in which the injection was made, S1 coded as 1, M1 coded as 0). The two injections with an injection volume of 0.5 µl were excluded here to reduce the influence of differences between the injected volumes.

The topography was analyzed by correlating the positions of the cells in different thalamic nuclei in mediolateral, dorsoventral and rostrocaudal direction (main axis directions) with the position of the injection sites in S1 or in M1. The positions of the injection sites were calculated for different directions (defined by the angle φ; φ between 0° and 359° in 1° steps) (Supp. Fig. 9). The rostral direction was represented by φ = 0° and the direction to the right was represented by φ = 90° (by definition). The position of the injection sites in S1 or in M1 was then calculated as:

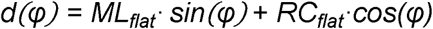

where *d* is the position of the injection site for the direction defined by φ, ML_flat_ is the mediolateral position of the injection site and RC_flat_ is the rostrocaudal position of the injection site (ML_flat_ and RC_flat_ were defined in relation to the two-dimensional representation of the cortex). The direction (φ) for a specific thalamic nucleus, a specific cortical area and a specific main axis direction, for which the corresponding correlation coefficient was highest, was defined as φ_max_.

Coordinates of the centroids of the cell positions (used for the analysis of the topography and the analysis of the differential distribution of cells in the thalamic nuclei) were defined by calculating the median of the positions of all cells detected in this nucleus in mediolateral, dorsoventral or rostrocaudal direction (e.g., the mediolateral coordinate of the centroid is defined as the median of the mediolateral coordinates of the detected cells). To reduce the influence of outliers (e.g., due to false positive detected cells), a centroid was only used for the analysis if at least eight cells had been detected in the respective thalamic nucleus.

The dice coefficient calculated for the cells that had taken up both fluorescent dyes was calculated by the following formula:

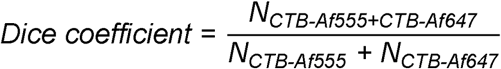

(*N_CTB-Af555+CTB-Af647_*: number of cells that had taken up CTB-Af555 and CTB-Af647, *N_CTB-Af555_* / *N_CTB-Af647_*: cells that had only taken up CTB-Af555 or CTB-Af647, respectively). Where possible, parametric tests were used for group comparisons; otherwise, non-parametric tests were applied. When comparing groups for the experiments with co-injections (Fig. 8), the Friedman test was used to compare the groups (Friedman, 1937). It should be noted that the statistical power of this test is limited when the sample is small, as in this case (Skillings & Mack, 1981). A p-value of 0.05 was used to assess statistical significance. Due to the large number of tests performed, the significance level was corrected using the Bonferroni method in the topographic analyses (Fig. 6, Supp. Tab. 4).

### Data and code availability

Data can be made available upon reasonable request. Code for data analysis (including the cell detection and atlas registration) is available at the following link: https://github.com/andreas-huth/thalamocortical-projections.

## Acknowledgements

We thank Dr. Rebecca Wallrafen for initial contributions, training in methods and supervision, and Dr. Carlo Beretta for contributing the idea of using StarDist to detect cell nuclei as seed points and for performing the training of the StarDist model. We want to thank Marion Schmitt and Claudia Koksch for technical support. We thank Dr. Nina Ullrich and Dr. Ivo Sonntag for training in methods. We gratefully acknowledge the advice and input that were given by Prof. Dr. Christoph Körber and Dr. Dr. Varun Venkateramani. We appreciate that Dr. Jan Meis from the Institute of Medical Biometry (Heidelberg University, Germany) gave helpful statistical advice. Finally, we thank Rainer Huth for the help with editing the manuscript.

---

**Supplementary Figure 1. Channel-specific detection of the fluorescent dyes.** (a) Example image for the CTB-Af488 channel (yellow), CTB-Af555 channel (green) und CTB-Af647 channel (red) for one animal with previous injection of CTB-Af488 in the primary somatosensory cortex. The relative signal intensity was determined by comparing the signal of a labeled cell (cyan in the left image) with the adjacent background signal (magenta in the left image). For the same areas, corresponding values were also determined in the microscopic images of the other two channels (middle and right images). Scale bar: 100 µm. (b) Relative signal intensities for a CTB-Af488 injection (b1), CTB-Af555 injection (b2) and CTB-Af647 injection. An increased relative signal intensity was found only for the channel for which it was to be expected (one-sided one-sample t-test, compared with the value 1.0). *p* ≥ 0.05 (ns), *p* < 0.01 (**), *p* < 0.001 (***). Dashed line at 1.0. Mean ± Standard deviation.

**Supplementary Figure 2. Adaption of the injection sites to the reference atlas.** (a) Creation of the two-dimensional representation of the cortex (coronal view). The distance between the longitudinal fissure and the boundaries of the cortical areas is measured (green line). S2: secondary somatosensory cortex (other abbreviations see Fig. 1). (b) Cerebral cortex in a flat representation. Grid space: 1 mm (abbreviations see Fig. 1). (c) Reference points are defined on the right hemisphere for the bright field image. (d) Corresponding reference points (same color as in c for corresponding points) are defined on the reference atlas slice. (e) Resulting deformation grid. (f) Injection site of a CTB-Af555 injection (green) after adaption to the reference atlas. Scale bars: 1 mm.

**Supplementary Figure 3. Detection of cell nuclei.** Top: Cell nuclei in the DAPI-channel images (left) were manually segmented with Labkit (right) to create the training data for StarDist. Middle: The cell nuclei are automatically segmented by the StarDist model (red). Scale bar: 50 µm. Bottom: The predictions of the model (red dots) were compared with manual annotations (orange crosses) (left image). Predicted cell positions were classified as false positive (fp, yellow), false negative (fn, cyan) or true positive (tp) (performed for *n* = 4 images). Most true positive detected nuclei were detected only a single time (tp-s), but a few nuclei were detected multiple times (tp-m) due to oversegmentation of the cell nuclei. The number of detected cells for each category is depicted on the right side for the four different images. Lines mark the mean. Scale bar: 100 µm.

**Supplementary Figure 4. Classification by Convolutional Neural Networks (CNNs).** (a) Training of three different CNNs (one example image, CTB-Af555 channel). Positions for labeled cells (blue crosses) and unlabeled cells (red crosses) have been marked. The positions were used to extract image crops of different sizes (the six smaller images are the extracted images belonging to the position marked with an arrow). Scale bar: 100 µm (larger images) and 10 µm (smaller images). (b) Accuracy of the CNNs for the three different models per channel (b1: CTB-Af488 channel, b2: CTB-Af555 channel, b3: CTB-Af647 channel) for the training data and the test data. (c) Confusion matrices for the different models (c1: CTB-Af488 channel, c2: CTB-Af555 channel, c3: CTB-Af647 channel; CNN1: 60 x 60 pixel, CNN2: 90 x 90 pixel, CNN3: 120 x 120 pixel).

**Supplementary Figure 5. Procedure of the adaption to the reference atlas.** Example for one brain slice (CTB-Af555 channel). Top: Adaption to the overview images. Three pairs of reference points (orange crosses) are defined on the average intensity projection of the LSCM stack (a1, a2) and the overview images (widefield fluorescence microscopic images) (a3, a4, a5). So, the cell positions (blue star symbols) could be transformed to the overview images. Bottom: Adaption to the reference atlas. Cell positions are then transformed from the overview images (b1, b2, b3) to the reference atlas (b4, b5, b6) by the definition of pairs of reference points (gold crosses). Scale bar: 0.5 mm (a1, a2, a3, a4, b2, b5) and 2 mm (a5, b1, b3, b4, b6).

**Supplementary Figure 6. Validation of the transformation from the LSCM images to the widefield fluorescence microscope images (overview images).** Two examples (a, b) for the transformation to the overview image. Blood vessels were segmented (blue) on the average intensity projection of the confocal stack (a1, b1; CTB-Af555 channel). The boundary points were then transformed to the overview images (a2, b2; CTB-Af555 channel). The transformed boundary line matched well with the boundary of the blood vessels in the overview image. Scale bars: 100 µm.

**Supplementary Figure 7. Calculation of the volume covered by the LSCM images**. The boundaries of the average intensity projection of the LSCM images (a, CTB-Af555-channel in green) were defined manually (red) and transformed to the overview images (widefield fluorescence microscope images) (b) and to the reference atlas (c). (d) Volume covered by the LSCM stacks as a fraction of the total volume for each thalamic nucleus (shown for all 16 animals). Dashed line at 0.5. Scale bars: 1 mm.

**Supplementary Figure 8: Number of detected cells per volume.** Shown for *n* = 19 injections in the primary somatosensory cortex (S1) and *n* = 13 injections in the primary motor cortex (M1) for the three used fluorescent dyes. Lines mark the median.

**Supplementary Figure 9: Calculation of the position *d*_i_ of the injection sites.** The direction for which the position of the injection site is calculated is defined by the angle φ (here: φ = 60°, shown are the injection sites in the primary somatosensory cortex (S1)). Injection sites displayed as circles (yellow: CTB-Af488, green: CTB-Af555, red: CTB-Af647). The calculation of *d*_i_ is based on the mediolateral position (*x*_i_) and the rostrocaudal position (*y*_i_) of the injection sites (regarding the two-dimensional representation of the cortex). Grid spacing: 1 mm.

**Supplementary Figure 10. Neurons projecting simultaneously to the primary somatosensory cortex and the motor cortex.** (a) Frontal view. (b) Top view. (c) Lateral view. Each point represents one detected cell. Left: Colors represent the respective animal. Right: Colors represent the respective thalamic nucleus to which the cells have been assigned to. ML: mediolateral, RC: rostrocaudal, DV: dorsoventral.

**Supplementary Table 1:**
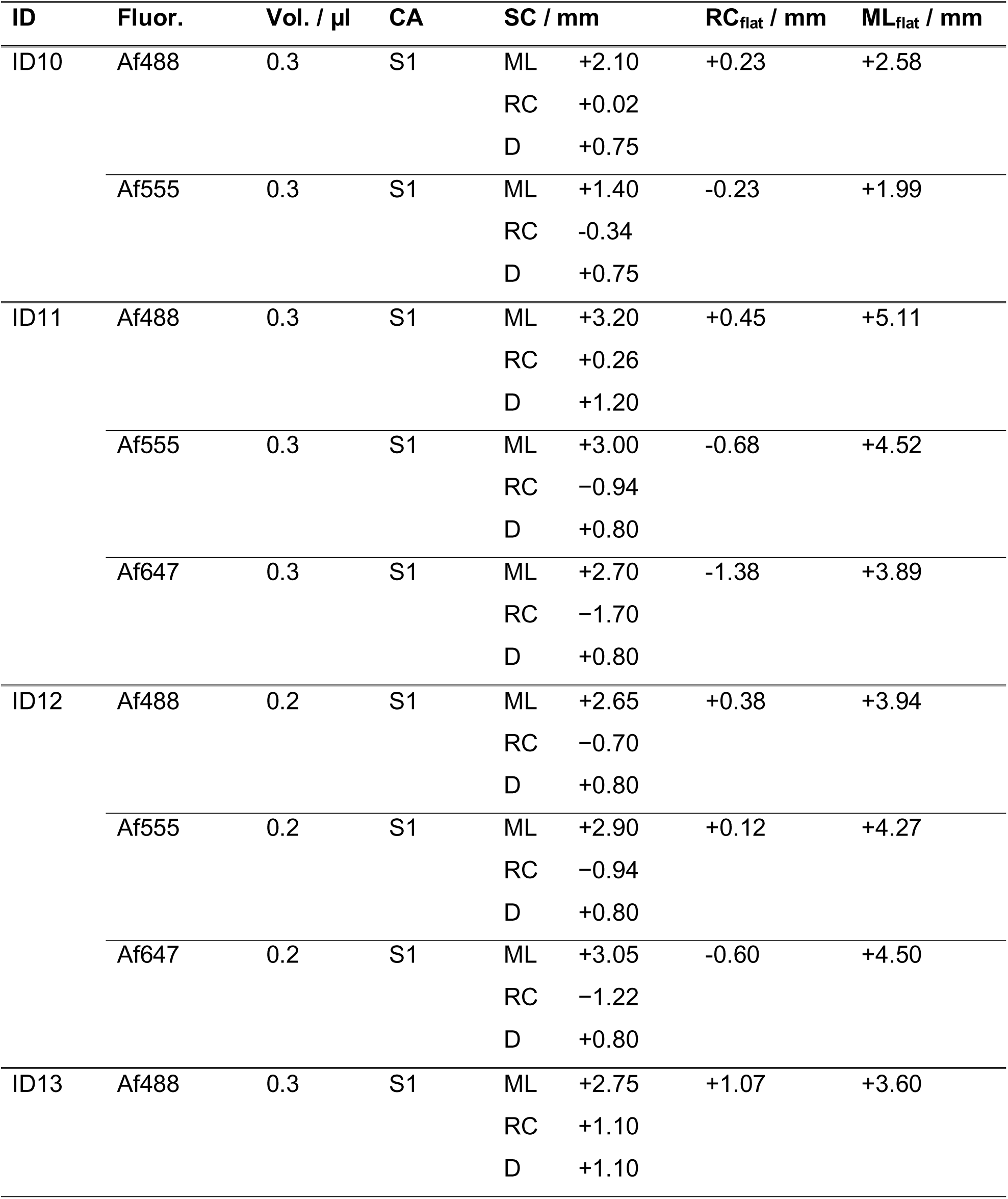
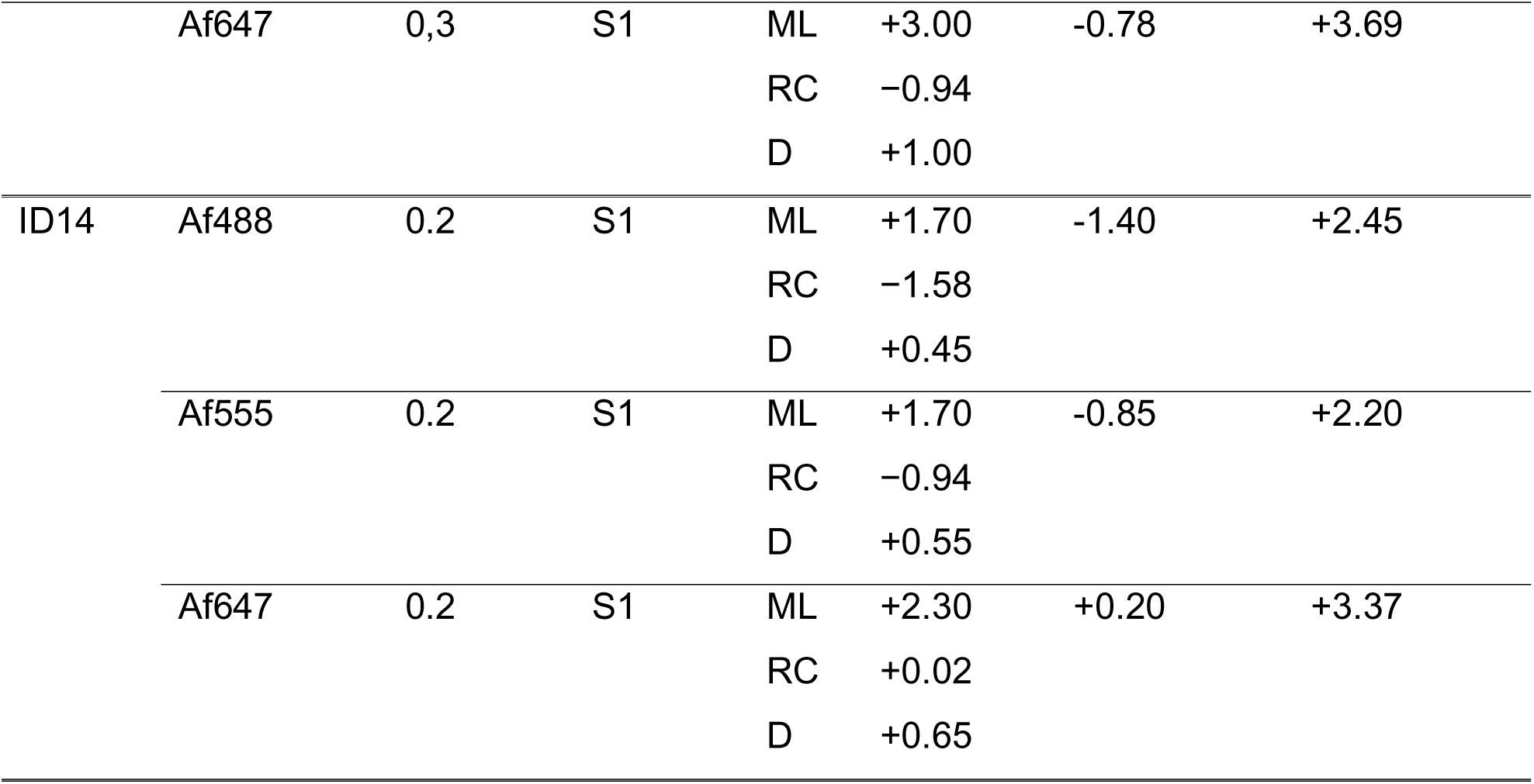
Injection sites (experimental group: injections only in S1). ID: animal identifier, Fluor.: fluorophore used for the injection (conjugated to CTB, Af488: Alexa Fluor 488, Af555: Alexa Fluor 555, Af647: Alexa Fluor 647), Vol.: injected volume, CA: cortical area of the injection, SC: used coordinates for the stereotactic injection (mediolateral (ML) and rostrocaudal (RC) position in relation to bregma, depth (D) in relation to the surface of the cortex / dura), RC_flat_: rostrocaudal position (in relation to the flat representation of the cortex), ML_flat_: mediolateral position (in relation to the flat representation of the cortex).

**Supplementary Table 2:**
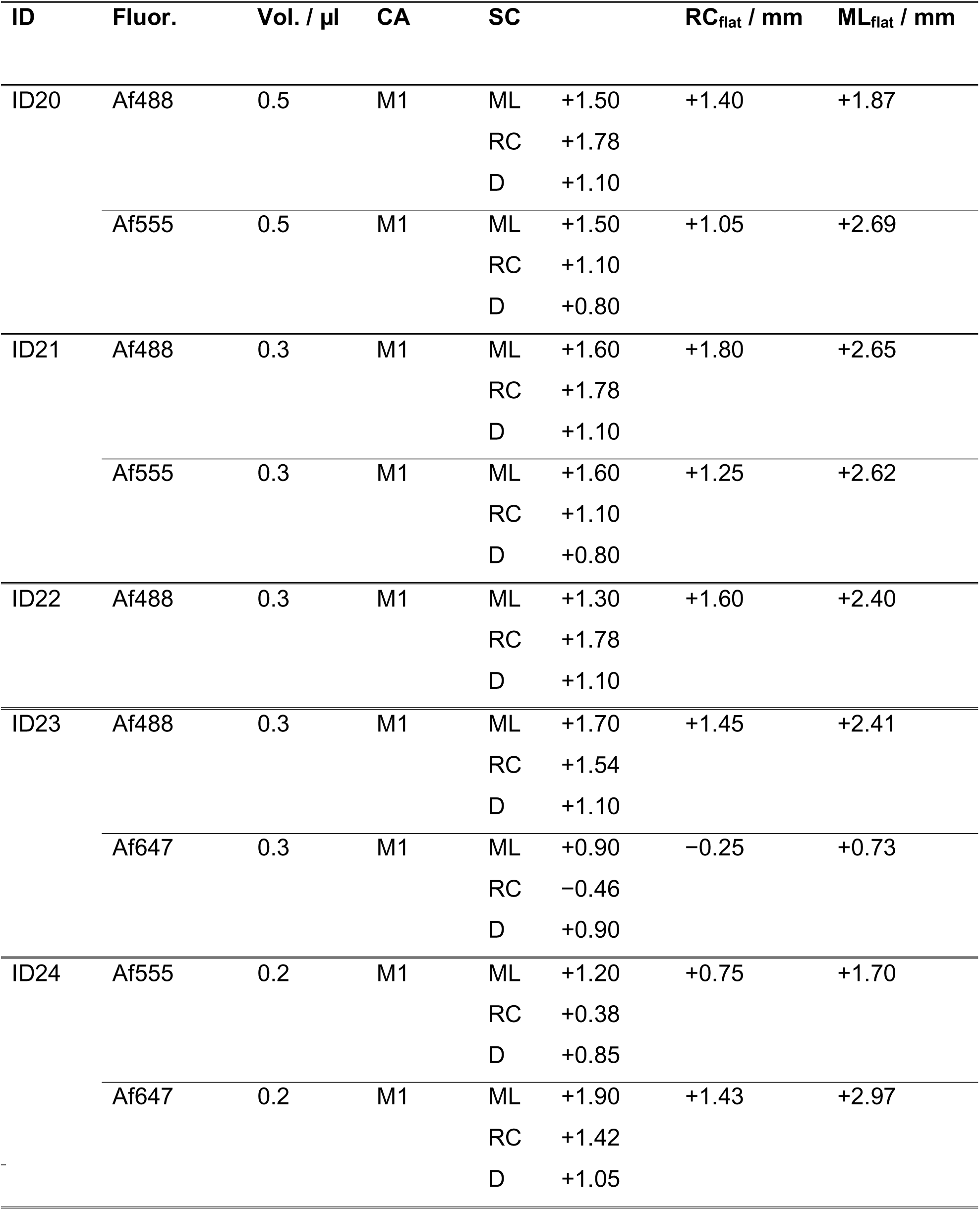
Injection sites (experimental group: injections only in M). ID: animal identifier, Fluor.: for the injection used fluorophore (conjugated to CTB, Af488: Alexa Fluor 488, Af555: Alexa Fluor 555, Af647: Alexa Fluor 647), Vol.: injected volume, CA: cortical area of the injection, SC: used coordinates for the stereotactic injection (mediolateral (ML) and rostrocaudal (RC) position in relation to bregma, depth (D) in relation to the surface of the cortex / dura), RC_flat_: rostrocaudal position (in relation to the flat representation of the cortex); ML_flat_: mediolateral position (in relation to the flat representation of the cortex).

**Supplementary Table 3:**
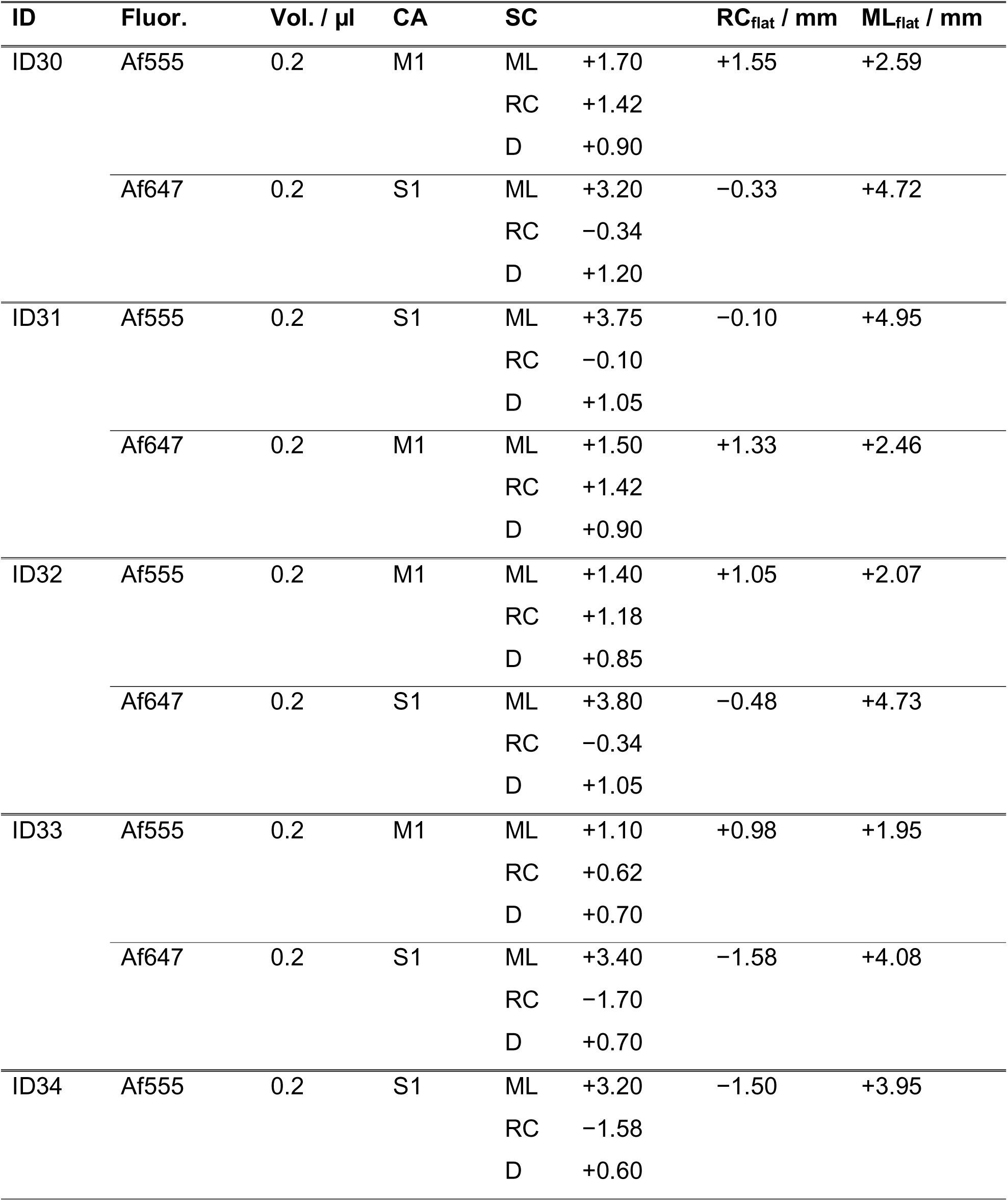
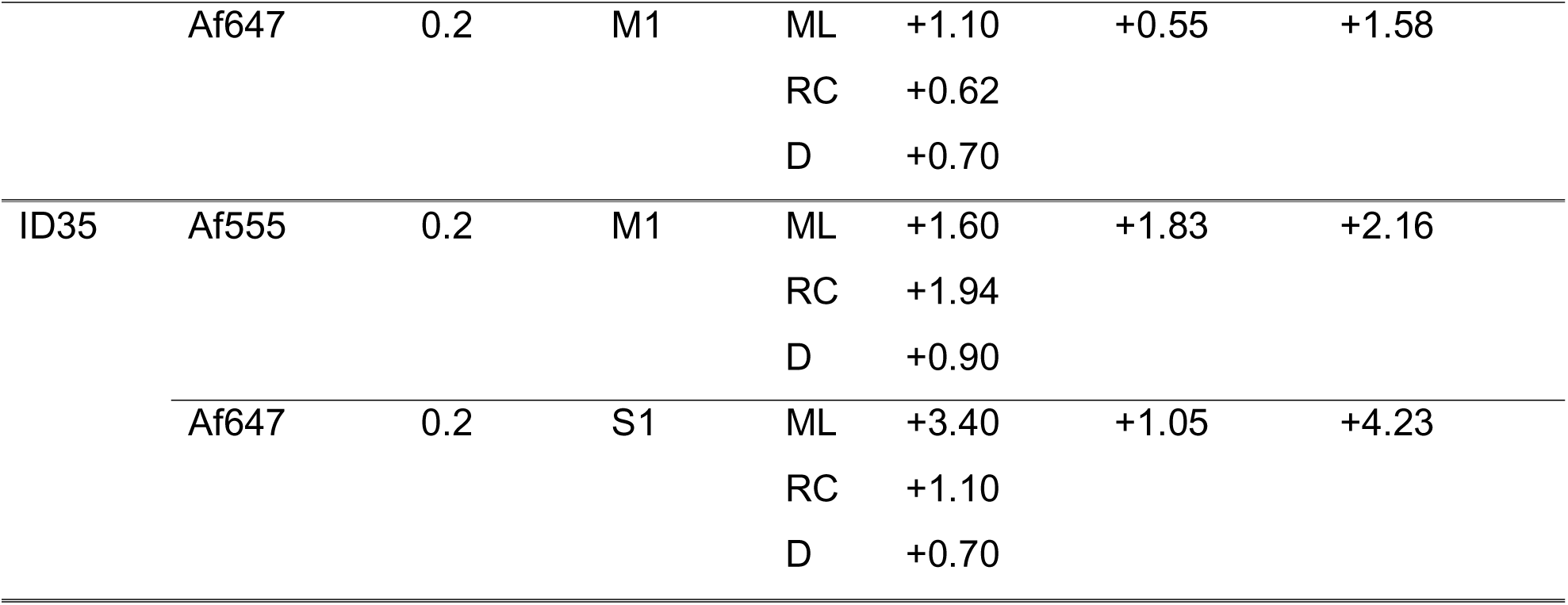
Injection sites (experimental group: co-injections). ID: animal identifier, Fluor.: for the injection used fluorophore (conjugated to CTB, Af488: Alexa Fluor 488, Af555: Alexa Fluor 555, Af647: Alexa Fluor 647), Vol.: injected volume, CA: cortical area of the injection, SC: used coordinates for the stereotactic injection (mediolateral (ML) and rostrocaudal (RC) position in relation to bregma, depth (D) in relation to the surface of the cortex / dura), RC_flat_: rostrocaudal position (in relation to the flat representation of the cortex), ML_flat_: mediolateral position in relation to the flat representation of the cortex).

**Supplementary Table 4:**
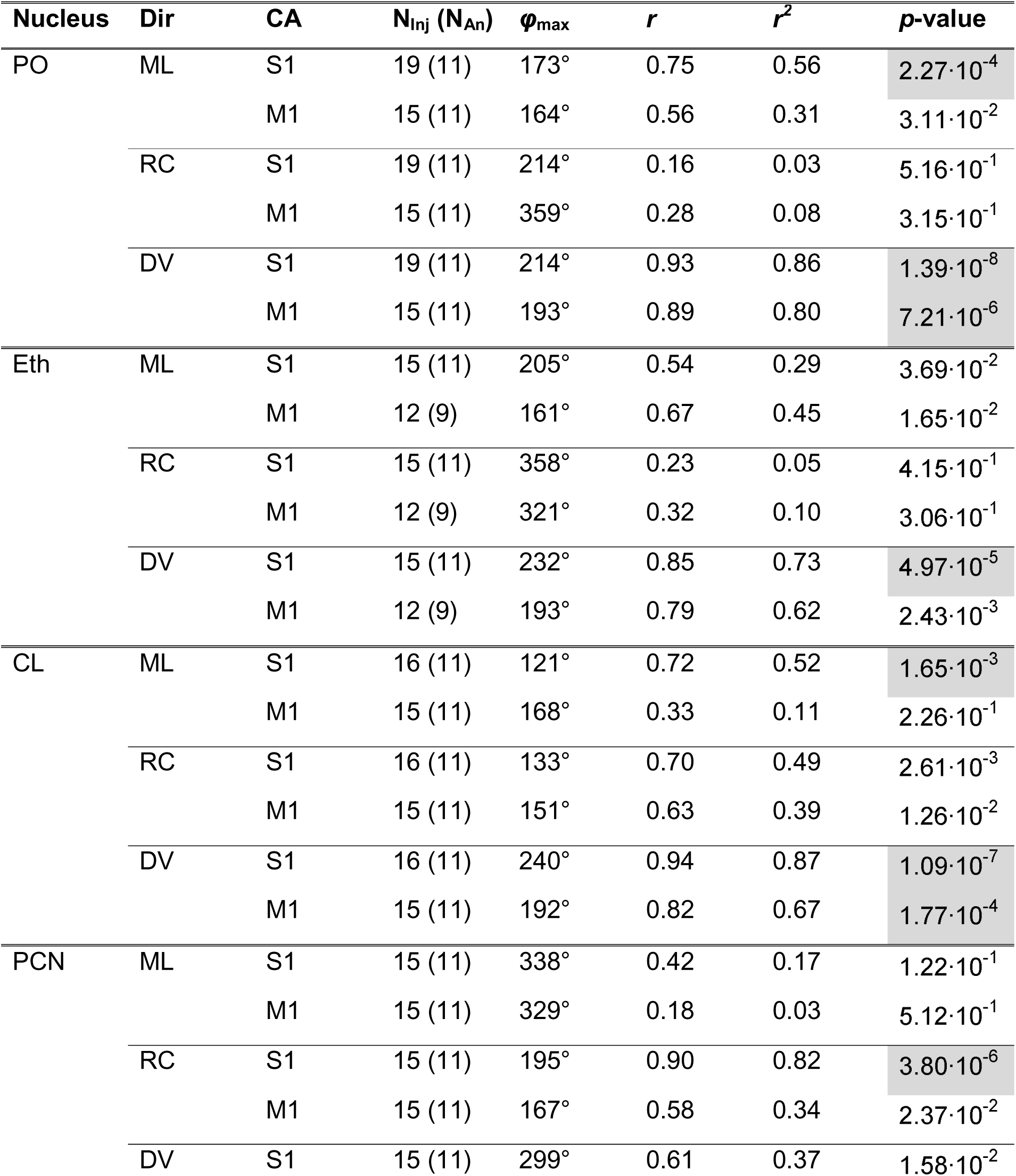
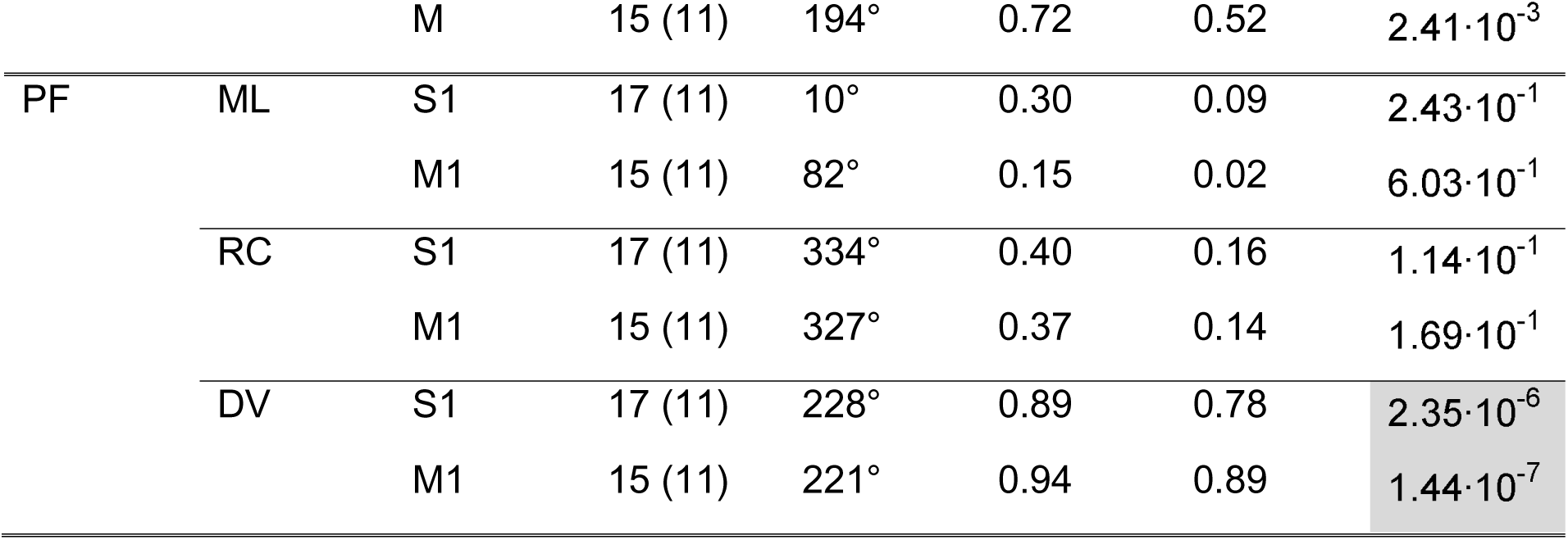
Topographical organization in different thalamic nuclei. Nucleus: thalamic nucleus, Dir: direction for the thalamic nucleus for which the topography was tested (ML: mediolateral, RC: rostrocaudal, DV: dorsoventral), CA: Cortical area to which the test for topographical organization refers to, N_Inj_: number of injections that were used for the analysis and number of different animals used (N_An_) (in brackets), φ_max_: direction for which *r* was maximal, *r*: Pearson correlation coefficient for φ_max_, *r^2^*: coefficient of determination for φ_max_, *p*-value: *p*-value for the linear regression analysis for φ_max_ (*p*-values below the corrected significance level (Bonferroni correction) are highlighted in grey).

